# Structures of ABCG2 under turnover conditions reveal a key step in drug transport mechanism

**DOI:** 10.1101/2021.03.03.433600

**Authors:** Qin Yu, Dongchun Ni, Julia Kowal, Ioannis Manolaridis, Scott M. Jackson, Henning Stahlberg, Kaspar P. Locher

## Abstract

ABCG2 is a multidrug transporter expressed widely in the human body. Its physiological substrates include steroid derivatives and uric acid. In addition, it extrudes many structurally diverse cytotoxic drugs from various cells, thus affecting drug pharmacokinetics and contributing to multidrug resistance of cancer cells. Previous studies have revealed structures of ABCG2 bound to transport substrates, nucleotides, small-molecule inhibitors and inhibitory antibodies. However, the transport mechanism is not well-understood because all previous structures described trapped states, where the reaction cycle was halted by the absence of substrates or ATP, mutation of catalytic residues, or the presence of inhibitors. Here we present cryo-EM structures of nanodisc-reconstituted human ABCG2 under turnover conditions containing either the endogenous substrate estrone-3-sulfate or the exogenous substrate topotecan. We found two distinct conformational states in which both the transport substrates and ATP are bound. Whereas the state turnover-1 features more widely separated NBDs and an accessible cavity between the TMDs, turnover-2 features semi-closed NBDs and an almost fully occluded cavity between the TMDs. The transition from turnover-1 to turnover-2 includes conformational changes that link the binding of ATP by the NBDs to the closing of the cytoplasmic side of the TMDs. The size of the substrate appears to control which turnover state corresponds to the main state in the transport cycle. The transition from turnover-1 to turnover-2 is the likely bottleneck or rate-limiting step of the reaction cycle, where the discrimination of substrates and inhibitors occurs. Our results provide a structural basis of substrate specificity of ABCG2 and provide key insight to understand the transport cycle.

## Introduction

ABCG2 is an ABC transporter of broad substrate specificity expressed in various tissues and tissue barriers^1-4^. Among its endogenous substrates are steroids, including estrone-3-sulfate (E_1_S), and the transporter has been reported to contribute to renal excretion of uric acid^5^. ABCG2 also functions as a multidrug transporter and exports a wide range of xenobiotics and pharmaceuticals, thus strongly impacting their pharmacokinetics^1-3^. Genetic polymorphisms of ABCG2 can lead to its dysfunction, which is associated with hyperuricemia and hypertension in humans^6^. Due to its broad substrate specificity, ABCG2 contributes to multidrug resistance (MDR) in cancer ^7,8^. For example, ovarian tumor and medulloblastoma cells over-expressing ABCG2 have shown increased resistance to chemotherapeutic agent including topotecan or mitoxantrone^9,10^. Considerable efforts have been directed at developing specific inhibitors of ABCG2 to counteract its activity in protecting tumour cells from anti-cancer drugs^11-17^. At present, the clinical applicability of such compounds is limited by their insufficient specificity, toxicity, or poor oral availability^18^. A detailed understanding of the mechanism of ABCG2 is therefore essential to advance the development of small-molecule modulators or inhibitors.

A series of initial structures have revealed the architecture of human ABCG2 and have provided insight into its binding of endogenous substrates as well as small-molecule inhibitors^19-21^. In the inward-open conformation, the pair of transmembrane domains (TMDs) form a slit-like cavity (cavity 1) that serves as a substrate binding pocket. Two phenylalanine side chains, one from each ABCG2 monomer, clamp substrates between their phenyl rings^21-24^. This can rationalize that ABCG2 substrates are generally flat, polycyclic, and hydrophobic compounds^20^. Inhibitors can also bind in cavity 1, both competing with substates for the binding pocket and interfering with the closing of the TMD dimer interface during the transport cycle^19-21^. Recent structural studies revealed that exogenous substrates (cytotoxic drugs used to treat various cancers) bind as single copies at the same general location as the endogenous substrate E_1_S^22,23^. In addition to inward-open structures, the structure of a variant with reduced catalytic activity, ABCG2_E211Q_, revealed an ATP-bound conformation with a closed nucleotide binding domain (NBD) dimer and a collapsed translocation pathway. This was taken as evidence supporting an ATP-driven, peristaltic extrusion mechanism^21^.

The common denominator of all published ABCG2 structures is that they represented trapped states determined under conditions where the transporter was prevented from cycling through its catalytically relevant conformations. This was accomplished either by the removal of ATP or transport substrates or by the addition of small-molecule inhibitors or externally binding inhibitory Fab fragments such as 5D3-Fab^19,25^. The strategy of locking intermediate states or reducing conformational flexibility has been at the heart of most high-resolution structural studies of multidrug ABC transporters since the publication of the Sav1866 structure^26^. Mutation of the catalytically essential Walker-B glutamate has often been used to trap ATP-bound states, where the nucleotide is sandwiched between the NBDs^21,27^. Occasionally, disulfide cross-linking was employed to trap intermediate states^28,29^. Such approaches were essential when using X-ray crystallography as a structural technique, since excessive conformational dynamics interfere with the generation of well-ordered 3D crystals. On the downside, these conditions differed from the physiological environment, where substrates, ATP, ADP, and Mg^2+^ are all present. As a result, the catalytic cycle of ABCG2 and other multidrug transporters is insufficiently understood.

While conformational rigidity is also beneficial for cryo-EM approaches, the single particle nature of cryo-EM studies provides an opportunity to investigate multidrug ABC transporters under turnover conditions, where distinct conformations are allowed to co-exist^30^. This approach is essential for understanding the transport mechanism because it reduces the chance of visualizing physiologically irrelevant conformational states or intermediates. In the present study we reconstituted human ABCG2 in lipid nanodiscs, which provide a near-native environment containing phospholipids and cholesterol. We applied turnover conditions using two distinct substrates and found a single major turnover state for the endogenous steroid E_1_S and two states for the larger, exogenous drug topotecan. Our study reveals key conformational changes that are essential for substrate recognition and translocation and allow ABCG2 to distinguish substrates from inhibitors.

## Results

### Structures of ABCG2 under turnover conditions

We determined the ATPase activity of ABCG2 reconstituted in nanodiscs (Supplementary Fig.1a) in the presence of E_1_S or topotecan (Fig.1a). To ascertain that the two substrates bound to nanodisc-reconstituted ABCG2, we measured the modulation of the ATPase rate compared to the absence of substrates. Compared to liposome-reconstituted ABCG2, the basal ATPase rate in nanodiscs is markedly elevated, as was observed previously^19-21^. This rate was further increased in the presence of E_1_S, but decreased in the presence of topotecan (Fig.1a). While the absolute values are higher than in liposomes, this observation, combined with the structural data, suggests that the two substrates indeed bound to nanodisc-reconstituted ABCG2. To mimic turnover conditions, we incubated ABCG2 with 5mM ATP, 0.5mM ADP, 5 mM MgCl_2_, and either 200 µM E_1_S or 100 µM topotecan, and applied the mixtures to EM grids. We collected single particle cryo-EM data of these two samples, which allowed us to determine three distinct high-resolution structures, one from the E_1_S-bound ABCG2 and two from topotecan-bound ABCG2. Both the NBDs and TMDs were well resolved in all three structures (Supplementary Fig.1b). Despite the fact that ABCG2 is a homodimer, we refined all maps in C1 symmetry and did not observe significant structural differences between the two monomers.

**Figure 1.**
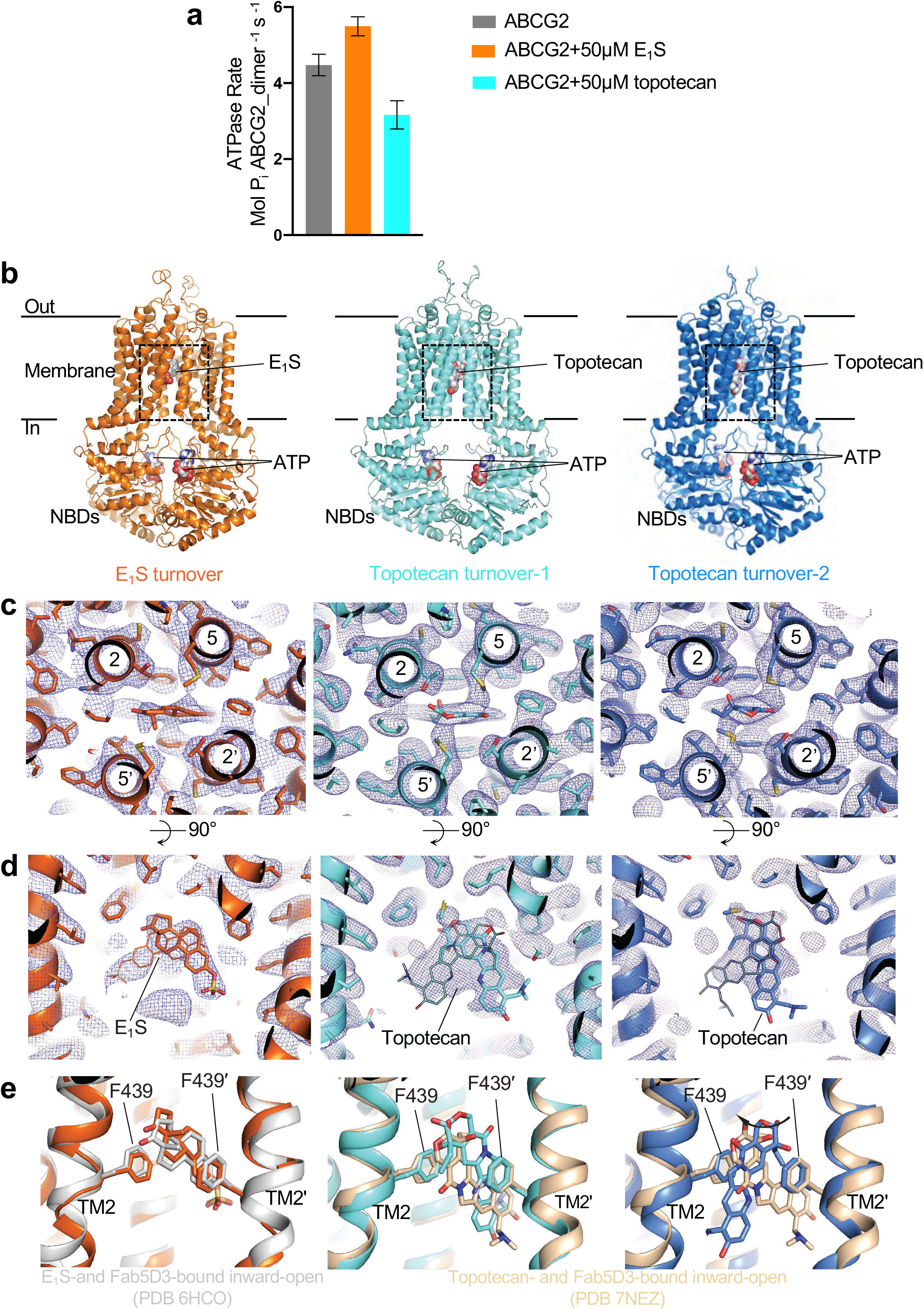
Structures of ABCG2 under turnover conditions. **a** ATPase activity of nanodiscs-reconstituted wild-type ABCG2 in the presence and absence of 50 μM E_1_S or 50 μM topotecan. The bars show means; error bars depict standard deviations (s.d.) (n=3, same batch of nanodiscs). **b** Ribbon diagrams of turnover structures. The colouring of the three structures is maintained throughout the figures and panels. Bound ATP, E_1_S and topotecan are shown in sphere representation and labeled. **c** Close-up view of substrate-binding pockets between the TMDs viewed from the extracellular side. TM helices are shown as ribbons and labelled, residues and bound substrates are shown as sticks. Substrates are at centre of the panels. Non-symmetrized EM density maps are shown as blue mesh. **d** Similar to **c**, but viewed from within the membrane. Bound substrates are labelled. Two possible orientations of bound substrates are shown for each density as thin or thick sticks, respectively. **e** Superposition of turnover states with structures of substrate-bound inward-open ABCG2 bound to 5D3-Fab fragments using the side chains of F439s as anchors. Left, E_1_S-bound ABCG2 structure (PDB 6HCO) is shown in grey cartoon. Middle and right, topotecan-bound ABCG2 structure (PDB 7NEZ) is shown in yellow cartoon. The substrates and side chains of F439 residues are shown as sticks.

The turnover sample containing E_1_S revealed a single, well-defined conformation resolved at 3.4Å resolution (Supplementary Fig. 2). In contrast, the turnover sample containing topotecan revealed two well-ordered structures, which we termed turnover-1 and turnover-2 states, resolved at 3.1Å and 3.4Å, respectively (Supplementary Fig. 3). The conformation of the E_1_S-bound structure was very similar to turnover-2 from the topotecan-containing sample (Supplementary table 1). We therefore refer to the E_1_S-bound structure as turnover-2 as well. The TMDs in the turnover states adopted inward-facing conformations but differed in their degree of opening towards the cytoplasmic side of the membrane. Turnover-1 is a more open conformation, whereas turnover-2 is a more closed conformation. All three structures revealed densities for transport substrates and two bound ATP molecules (Fig. 1b).

While we assume that a closed conformation similar to that observed in the structure of the ATP-bound ABCG2_*E211Q*_ variant is present under turnover condition^21^, we did not observe a defined class of such particles in our turnover samples. This suggests that this closed conformation is not a low-energy state of the wild-type protein under turnover conditions. We also did not observe a collapsed apo-state similar to that reported recently for apo-ABCG2^22^. In the topotecan-bound ABCG2 sample, the dominant class (88% of the ordered particles) belonged to the turnover-1 state, whereas 12% particles belonged to the turnover-2 state (Supplementary Fig. 3). This suggests that turnover-1 is the lowest-energy state of ABCG2 in the presence of topotecan, whereas turnover-2 is the lowest-energy state in the presence of E_1_S. In all turnover structures, the leucine gate (formerly called leucine plug), which separates cavity 1 from cavity 2, is closed (Supplementary Figs. 4a, 4b).

### E_1_S and topotecan binding site

We initially expected that under turnover conditions, transport substrates would either be too mobile to be observed or not present at all. To our surprise, we found strong densities for bound substrates lodged in cavity 1 on the two-fold molecular symmetry axis of ABCG2 (Figs. 1c, 1d). Substrate recognition and binding are similar as in the previously reported, inward-open structures^21-23^. The polycyclic cores of E_1_S and topotecan are sandwiched between the F439 side chains of the two ABCG2 protomers (Figs. 1c, 1d). Only one substrate molecule can be fitted inside the density, as fitting two molecules would introduce serious steric clashes (Fig. 1d). The density suggests a certain degree of flexibility of bound substrates, in particular for topotecan (Fig. 1d). In the structure containing E_1_S (turnover-2), the substrates could be fit in two orientations related by a 180° rotation. Due to the C1 processing of the data, one orientation is more prominent. When compared to the previously reported structure of E_1_S-and 5D3-Fab-bound ABCG2 in an inward-open conformation, E_1_S appeared slightly shifted (∼1 Å) towards the leucine gate and thus towards the external side of the transporter (Fig. 1e). In the topotecan-containing turnover structures, the shape of the density feature covering the drug is different in turnover-1 versus turnover-2. In turnover-1, topotecan can be fitted in two orientations related by a 180° rotation (Fig. 1d middle), similar to what was observed in an earlier structure of topotecan- and 5D3-Fab-bound, inward-open ABCG2^23^, and with no significant shift of bound topotecan in the direction of the leucine gate. In turnover-2, the topotecan density was narrower in the horizontal dimension and longer in the vertical dimension, probably reflecting a rotation of the drug compared to the turnover-1 state (Fig. 1d right). The changes in EM density could be visualized by 3D variability analysis of the final particles using the program cryoSPARC (Suppl. movies 1 and 2)^31,32^.

### Coupled NBD and TMD motions

ABC transporters use the energy of ATP binding and hydrolysis to translocate substrates across the membrane via an alternating access mechanism^33-35^. In the course of the ATPase cycle, the NBDs cycle between closed and open conformations to hydrolyze ATP or to release ADP and phosphate. This drives conformational changes in the TMDs. The separation of the NBDs in our turnover-1 and turnover-2 structures is smaller than that in the fully inward-open, topotecan-bound ABCG2 structure^23^, but larger than that in the ATP-bound structure of ABCG2_E211Q_ (Fig. 2a)^21^. A comparison of these four structures shows that as the NBDs approach each other, so do the cytoplasmic sides of the TMDs (Fig. 2a). The TMDs rotate relative to the NBDs (Supplementary Fig. 4c). Compared to the inward-open conformation, access to cavity 1 from the inner leaflet of the membrane is reduced in turnover-1, as the gap between transmembrane helices TM1 and TM5’ of the opposing ABCG2 monomer is narrower (Figs. 2b, c). In turnover-2, the gap is even smaller, causing the entrance to cavity 1 from within the membrane to be completely sealed and even the opening to the cytosol to be reduced. As a result, a larger substrate such as topotecan could not enter into, or exit from, cavity 1 without a spreading of the cytoplasmic sides of the TMDs (Fig. 2c). The degree of NBD closing and TMD gap narrowing can be measured by comparing the distances of the coupling helices (CpH), which are located in the first intracellular loop of the TMDs and transmit conformational changes at the NBD-TMD interface. In turnover-1, the coupling helices are moved towards each other by ∼1Å compared to the fully inward-open, topotecan- and 5D3-Fab-bound structure (Fig. 2d). In turnover-2, the distance between the coupling helices is reduced by another 4.0 Å. The distances between the TMDs are shifted accordingly (Supplementary Fig. 5). This reveals a strict coupling between NBD and TMD closing.

**Figure 2.**
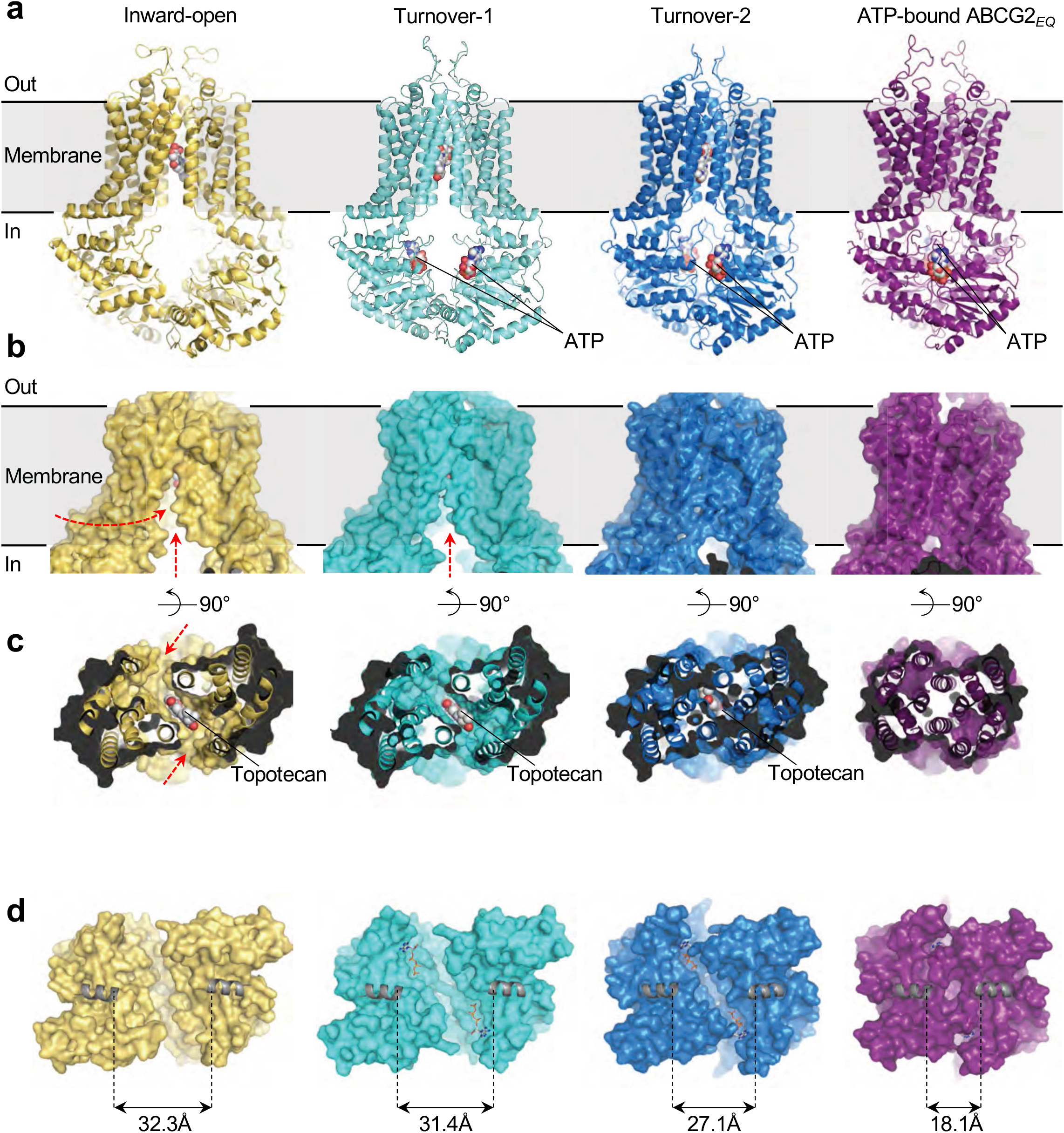
Coupled NBD and TMD closing. Structures of four states are shown: “Inward-open” depicts topotecan-and 5D3-Fab-bound ABCG2 (PDB ID 7NEZ), the turnover states are from this study, “ATP-bound ABCG2_E211Q_” depicts the closed conformation (PDB ID: 6HBU). **a** Ribbon diagrams with substrates and ATP shown as spheres. **b** Surface representations of ABCG2, emphasizing the distinct TMD conformations. Red arrows depict openings to cavity 1 from within the lipid bilayer or from the cytosol. **c** View of substrate-binding site in cavity 1 of ABCG2. The TMDs are shown in ribbon and surface representation and viewed from the cytoplasmic side of the membrane. Substrates are shown as spheres and labelled. Red arrows depict openings to cavity 1 from within the lipid bilayer. **d** Surface representations of the NBDs viewed from the membrane. Bound ATP is shown as sticks. The coupling helices (CpHs) are shown as grey cartoons. Distances between CpHs are indicated.

In addition to the motion of the NBDs as full domains, we observed a rotation of the RecA-like and the helical sub-domains relative to each other, which caused the helical sub-domain to further approach the opposite NBD. Superimposing the RecA-like domains and measuring the relative angles of the α-helical domains revealed rotations of 2°, 19°, and 24° between the fully inward-open conformation and turnover-1, turnover-2, and the ATP-bound closed state (Supplementary Fig. 6).

A comparison of the topotecan-bound turnover-1 and turnover-2 structures revealed a key NBD segment contributing to functionally relevant NBD-NBD and NDB-TMD domain contacts. As ABCG2 transitions from turnover-1 to turnover-2, the segment between Q181 and V186 undergoes a conformational change and facilitates key inter-domain contacts (Figs. 3a, b). V186 is the first residue of the VSGGE motif (equivalent to LSGGQ more commonly found in ABC transporters)^36^. This motif is among the most conserved in ABC transporters and pins bound nucleotides against the Walker-A and Walker-B motifs of the opposing NBD. We now found that in turnover-2, Q181 and F182 facilitated contacts between TM1 of one ABCG2 monomer and TM5’ of the other (Fig. 3a, b), thus helping stabilize the TMD domain closure. In addition, the arginine residue R184 provides a direct contact, in the form of a cation-π interaction, with the adenine moiety of the ATP molecule bound to the opposite NBD (Fig. 3d). Neither of these contacts are formed in the turnover-1 structure (Fig. 3c).

**Figure 3.**
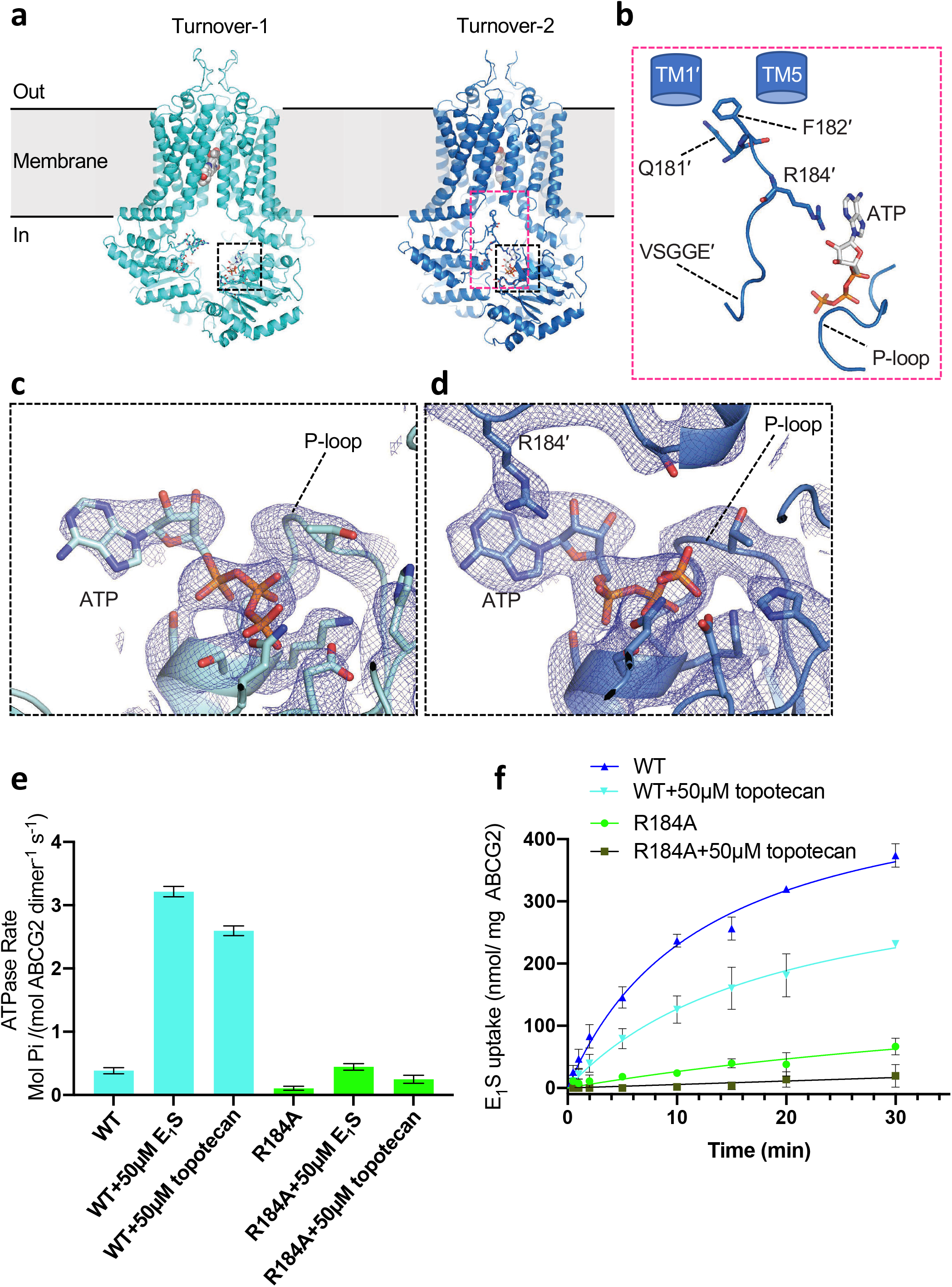
Domain interfaces and the role of R184. **a** Ribbon diagrams of topotecan-bound turnover states. ATP is shown as sticks, topotecan is shown as spheres. Dashed black and red boxes show domain interfaces with close-up views in panels **b**-**d. b** Close-up view of domain interface in turnover-2 conformation. TM1’ and TM5 are schematically shown as blue cylinders. P-loop, X-loop, and signature motif (VSGGE) are shown in cartoon mode. ATP and key side chains involved in surface interactions are shown as sticks. **c** and **d** Close-up views of ATP-binding sites of turnover-1 and turnover-2 states, as indicated in panel **a**. ABCG2 is shown as a ribbon diagram, key residues and bound ATP are shown as sticks and labeled. Non-symmetrized EM density maps are shown as blue mesh. **e** ATPase activity of liposome-reconstituted wild-type and mutant ABCG2 (R184A) in the presence and absence of 50μM E_1_S or 50 μM topotecan. Error bars depict s.d. (n=3, same batch of liposomes). **f** ABCG2-catalyzed E_1_S transport into proteoliposomes by wild-type ABCG2 or R184A variant in the presence or absence of 50 μM topotecan. Error bars depict s.d. (n=3).

We analysed the role of R184 by mutating it to an alanine (Supplementary Figs. 7a, 7b). We reconstituted the purified ABCG2_*R184A*_ variant in proteoliposomes and performed ATPase and transport assays (Figs. 3e, f). We found that in line with its location in the structure, both the ATPase and transport activities of ABCG2_*R184A*_ were greatly reduced. As was observed for WT ABCG2, the ATPase rate of ABCG2_*R184A*_ was stimulated by E_1_S and topotecan (Fig. 3e). Furthermore, the E_1_S transport rate was similarly reduced for WT ABCG2 and ABCG2_*R184A*_, suggesting that the mutation of R184 to alanine neither interfered with substrate binding nor abolished the NBD-TMD coupling. Given that the EC_50_ values of E_1_S- and topotecan-induced stimulation of the ATPase activity of ABCG2 were similar (15.7 and 11.7 µM, respectively, Supplementary table 2)^21,23^, the reduction of the initial E_1_S transport rate by ∼50% in the presence of topotecan is in line with the structural data in that topotecan and E_1_S bind and compete for the same binding pocket both in WT ABCG2 and in the ABCG2_R184A_ variant (Supplementary Fig 7c).

### The role of R482

The single-nucleotide polymorphisms R482G and R482T, originally cloned from drug-resistant cancer cell lines^37,38^, have been reported to alter the substrate specificity of ABCG2^39^. Cells expressing ABCG2_R482G_ were found to efficiently extrude rhodamine-123 or doxorubicin but were less resistant to topotecan, an anti-cancer drug that inhibits topoisomerase^7,38,40,41^. It was therefore suggested that R482 influenced substrate transport and ATP hydrolysis, but not substrate binding^42,43^. R482 is located in TM3 and does not directly contact bound drugs^21-23^. Rather, it contacts TM2 which contains the key residue F439 that interacts with substrates in cavity 1^19-21^. Intriguingly, we found that the side chain of R482 adopts distinct conformations in the turnover-1 and turnover-2 structures (Fig. 4). In turnover-1, the side chain points towards the cytoplasmic side of the membrane, where the guanidinium group forms a hydrogen bond with the side-chain of S443 of TM2 (Figs. 4b and 4c, left). In contrast, the R482 side chain is rotated towards the external side of the membrane in turnover-2, where the guanidinium group contacts side chains and main chain atoms from TM2 and TM4 in the direct vicinity of F439 (Figs. 4b and 4c, right). This suggests a structural role for R482 in facilitating the conformational changes required to transition from turnover-1 to the turnover-2 state (visualized in Suppl. movie 2). The impact of mutations of R482 on drug recognition is therefore likely of an allosteric nature.

**Figure 4.**
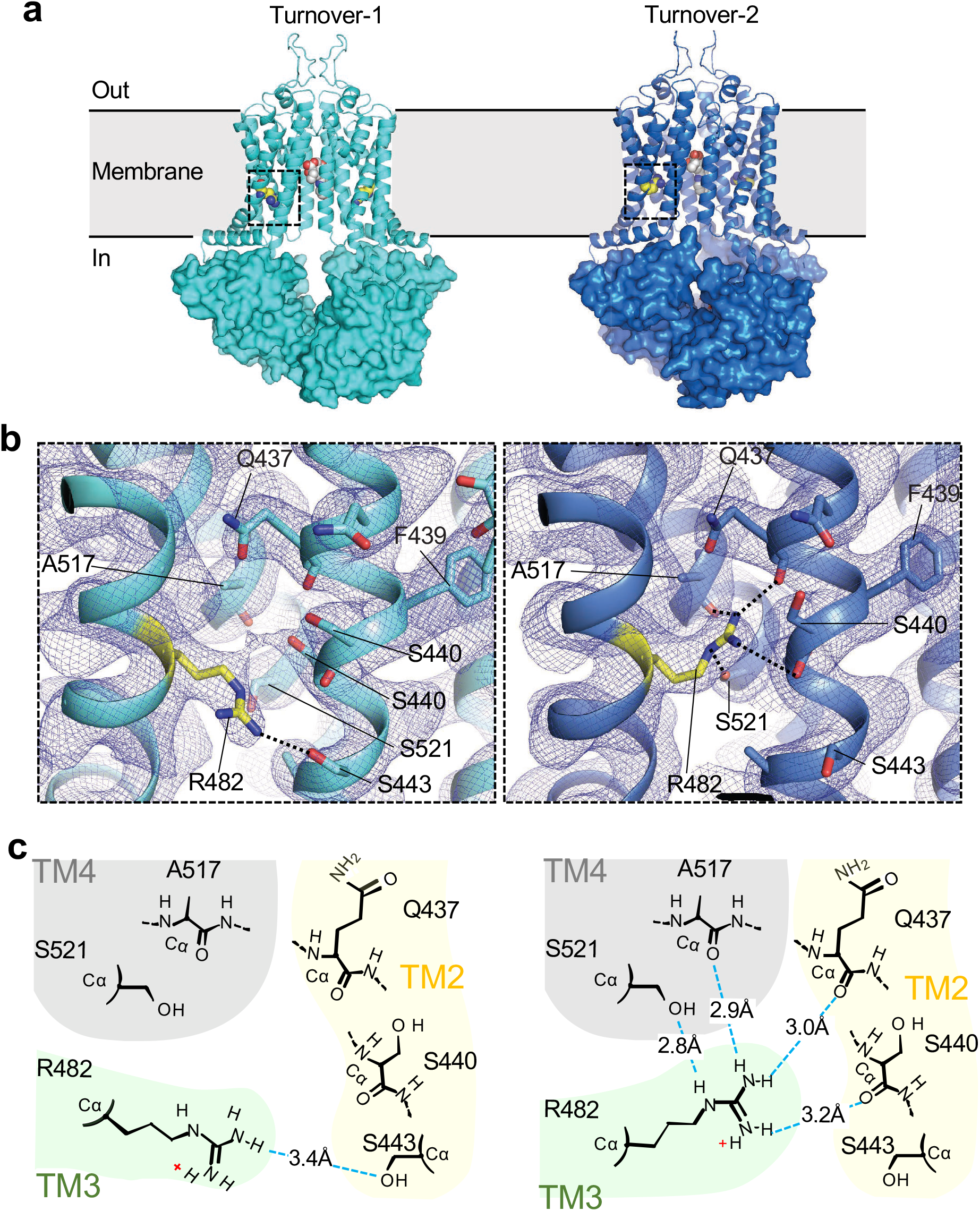
Conformational change of R482 in turnover states. **a** Ribbon diagrams of topotecan-bound turnover states, with NBDs shown as surfaces. Bound topotecan and the side chain of R482 are shown as spheres. Dashed black boxes show regions around R184. **b** Closeup views of TMD regions indicated in **a**. ABCG2 is shown as a ribbon diagram in turnover-1 (left) and turnover-2 (right) states. R482 and surrounding residues are shown as sticks and labelled. Non-symmetrized EM density is shown as blue mesh. Hydrogen bonds are indicated as black dashed lines. **c** Schematic diagram of hydrogen bonds formed by R482 in turnover-1 (left) and turnover-2 (right). H-bonds are shown as blue dashed lines. Yellow shading highlights residues of TM2, green shading highlights residues of TM3, grey shading highlight residues of TM4.

### Steric clashes of inhibitors in turnover-2 state

Our structures provide additional insight into the mechanism of small-molecule inhibitors of ABCG2. The reduced size of cavity 1 in the turnover-2 conformation would cause steric clashes with inhibitor molecules that can bind to inward-open ABCG2^20^. Fig. 5 reveals that whereas there is sufficient space in cavity 1 in the turnover-2 state for bound substrates E_1_S or topotecan, the inhibitors MZ29 and MB136 do not fit. The concerted motions of TM1, TM2 and TM5 occurring during the transition from turnover-1 to turnover-2 reduce the volume of cavity 1 by approximately one third, from ∼1300 Å^3^ in turnover-1 to ∼830 Å^3^ in turnover-2 ^44^, preventing the binding of the larger inhibitor molecules. We conclude that small-molecule inhibitors of ABCG2, once bound to the protein, block the transition from turnover-1 to turnover-2. Unlike in B-subfamily ABC transporters, this all but prevents the NBDs from forming a closed dimer as required for ATP hydrolysis, which can explain why ABCG2 has a strongly reduced ATPase rate in the presence of inhibitors^20^, whereas ABCB1 does not^20,29,45,46^.

**Figure 5.**
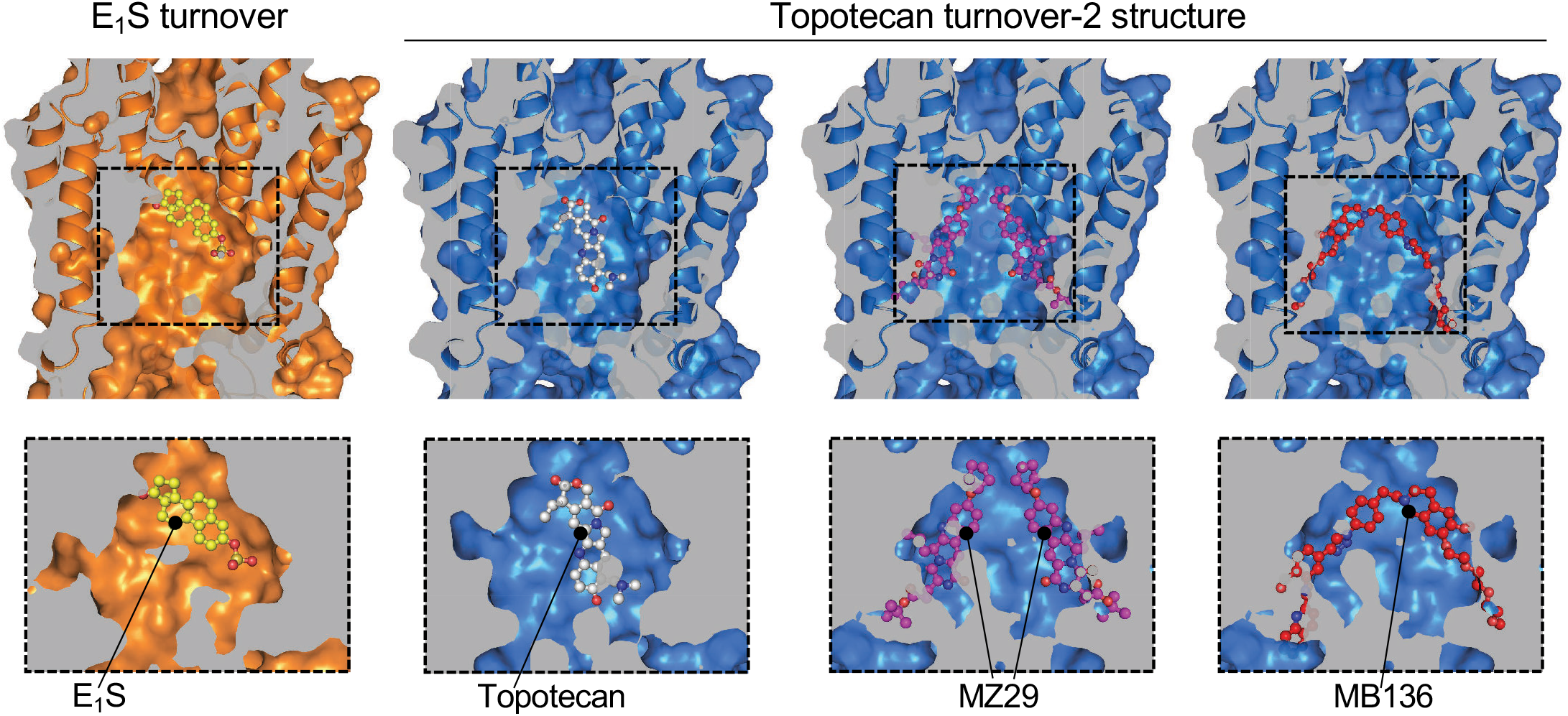
Cavity 1 in turnover-2 state accommodates substrates but not inhibitors. Vertically sliced views through E_1_S-bound (left) and topotecan-bound (right) turnover-2 structures. The top row of panels shows the TMDs both as ribbons and in surface representation. The bottom row shows close-up views of cavity 1, with ABCG2 in surface representation. Bound E_1_S and topotecan are shown in mixed stick/sphere representation as built in the structures. Inhibitor structures were extracted from inward-open ABCG2 structures containing these inhibitors (PDB 6ETI and 6FEQ, respectively) and manually placed into cavity 1 of the topotecan turnover-2 structure, emphasizing the resulting steric clashes.

## Discussion

In contrast to earlier structural studies of ABCG2, we could not predict whether the transporter would adopt a single conformation or multiple distinct conformations under turnover conditions. We also could not predict which state would correspond to the lowest energy state in the transport cycle. Our finding that two distinct inward-open conformations containing both a bound transport substrate and two ATP molecules corresponded to the lowest energy states was nevertheless unexpected. This arrangement was often considered to be of higher energy or even transient in ABC transporters, and thus expected to quickly convert to intermediates where either the transport substrate was absent, or ATP had been hydrolyzed^47,48^. Indeed, an analysis of bovine ABCC1, another multidrug resistance protein, captured a post-hydrolysis state under turnover conditions featuring a closed NBD dimer and a (slightly) outward-open substrate translocation pathway. This resembled a previously reported structure of a catalytically impaired ABCC1 variant containing a Walker-B glutamate to glutamine mutation^47^. In contrast to bovine ABCC1, a bacterial ABC transporter of peptides, TmrAB, was found to adopt multiple inward-open conformations under turnover conditions. These varied in how widely the NBDs were separated, the degree of inward opening of the translocation pathway, and whether bound nucleotides or substrates could be identified^48^. Our findings with human ABCG2 are distinct from both of those studies, which most likely reflects the mechanistic diversity within the ABC transporter superfamily^34^.

By combining previously reported structures of trapped states and our new turnover structures, we can now propose a structure-based hypothesis of the complete transport cycle of ABCG2 (Fig. 6). Only conformational states that are required for transport are included in this scheme. Our new turnover structures have a key role in this mechanistic scheme. Given that E_1_S did not promote the turnover-1 state, we concluded that ABCG2 can directly proceed to turnover-2 when smaller or endogenous substrates are bound. We anticipate a similar situation with urate, an even smaller molecule than E_1_S. For these substrates, turnover-2 appears to immediately precede the rate-limiting step of the transport cycle. In contrast, turnover-1 was the dominant class in the topotecan turnover sample. This suggests that the transition from turnover-1 to turnover-2 may be the step where exogenous compounds are “tested” for their suitability as transport substrates. Because topotecan is larger than E_1_S, the transition from turnover-1 to turnover-2 is slower. We conclude that the degree of TMD and NBD closing in the lowest energy intermediate correlates with the shape and size of ABCG2 substrates. More generally, compounds that bind in cavity 1 and allow ABCG2 to adopt the turnover-2 conformation may be transported, even if their transport rates are slower than that of E_1_S. In contrast, in the presence of strongly bound inhibitors such as Ko143 and its derivatives^13,20^, ABCG2 cannot advance to the turnover-2 conformation.

**Figure 6.**
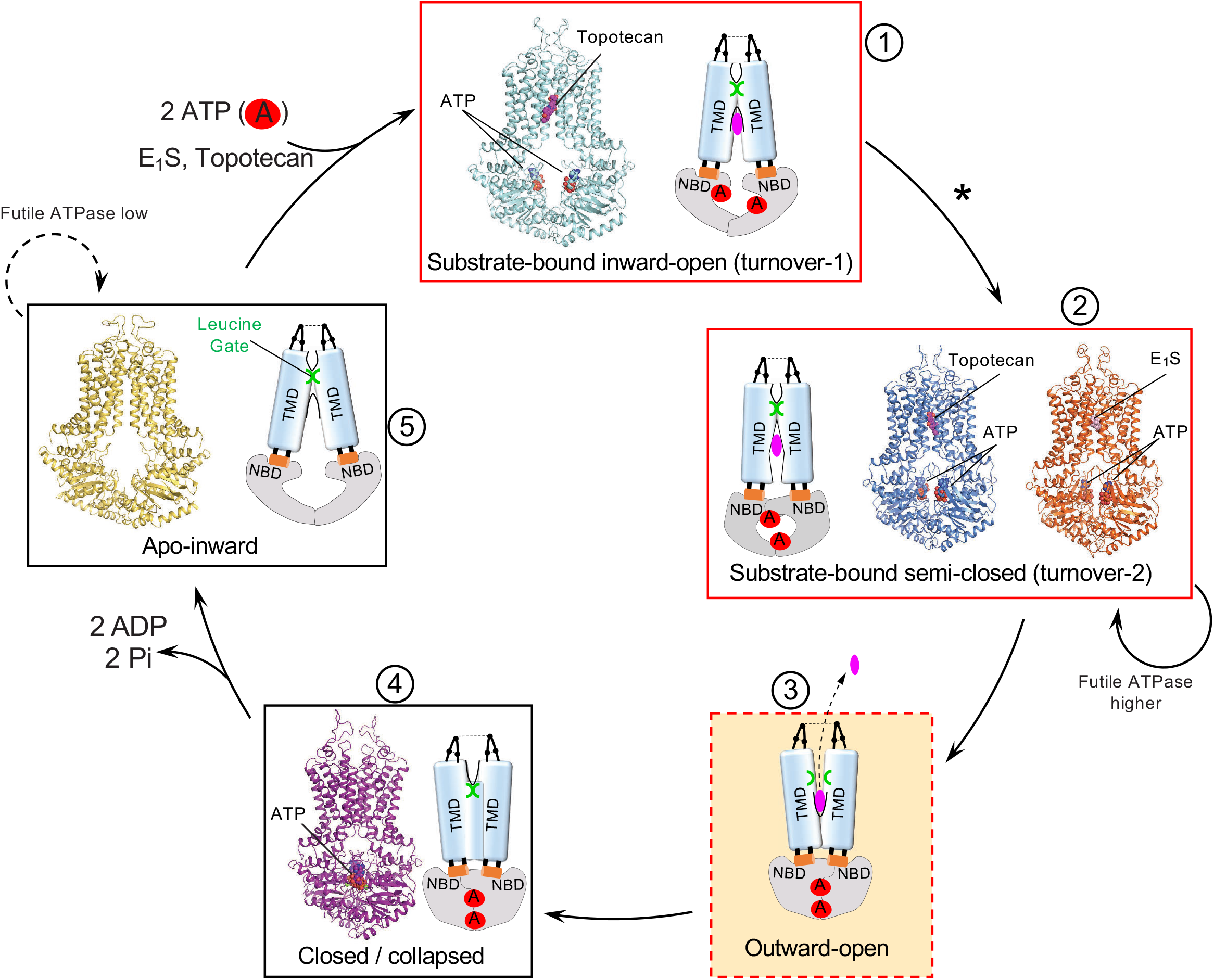
Transport mechanism of ABCG2. Schematic of structure-based, proposed transport cycle of ABCG2. Key states are shown as boxes and numbered. Where available, structures are shown as ribbon diagrams with bound nucleotides and substrates shown as spheres and labelled. Cartoon schematics of the relevant states are drawn to emphasize the conformations of the TMDs and NBDs, identify cavities, indicate the presence of bound drugs and as well as the state of the leucine gate. Previously determined structures correspond to PDB IDs 5NJ3 (apo-inward) and 6HBU (closed conformation). No structure is available of the proposed outward-open conformation (state 3).

Our results further assign a key role to the segment 181-184 of ABCG2. It was previously reported that the polypeptide segment leading up to the LSGGQ motif appeared to have a role in stabilizing TMD-TMD interfaces in B-subfamily ABC transporters, and the term X-loop was introduced to account for this role^26^. Rather than an aromatic residue to stabilize the adenine moiety of bound nucleotides, as observed in the A-loops of most ABC transporters^27,49-51^, ABCG2 relies on R184 from the opposite protomer to assume this function. R184 therefore appears to have a dual role in the turnover-2 state: It pioneers the contact between the two NBDs while simultaneously strengthening the binding of ATP through a cation-π interaction with the adenine moiety. The initial NBD-NBD contact is probably important because the signature motif is not yet close enough to interact with the ribose and triphosphate moieties of the ATP molecule.

Our mechanistic scheme indicates two distinct levels of ATPase activity, a basal and a stimulated level (Fig. 6). In the absence of substrates, ABCG2 displays a basal ATPase rate that generates 0.6 molecules phosphate per ABCG2 dimer and second. In the presence of E_1_S, this value increases to 2.3. We compared this value to the initial E_1_S transport rate, which is 0.1 molecules E_1_S per ABCG2 dimer and second. If ABCG2 hydrolyses two molecules of ATP in one productive transport cycle, the net futile ATPase rate in the presence of E_1_S is 1.5 molecules phosphate per ABCG2 dimer and second, which is 2.5 times higher than in the absence of E_1_S. We conclude that substrates not only increase the ATPase rate of ABCG2 due to productive transport, but also increase the basal / futile ATPase rate. This suggests that ABCG2 may require multiple attempts (each consuming ATP) before substrate extrusion is successfully accomplished.

In both turnover states described here, the leucine gates are closed. This is notable in the case of turnover-2, where not only the NBDs are semi-closed, but the cytoplasmic ends of the TMDs are also partially closed. This suggests that the opening of the leucine gate may only occur once the cytoplasmic ends of the TM helices are further pushed together and peristaltic pressure is exerted on bound substrate. We speculate that at this point, ABCG2 must adopt an outward-facing conformation, in which the leucine gates are opened and substrate can escape via cavity 2 into the extracellular environment. Such a conformation may be transient in nature given that we have not observed it in our turnover samples. Further studies are required to capture ABCG2 or an appropriate ABCG2 variant in an outward-open conformation.

In conclusion, our results provide key insight into the transport cycle of ABCG2 and help understand how small-molecular compounds act as substrates or inhibitors. Our study may also have predictive value for understanding the mechanisms of other ABC transporters, in particular of the G-subfamily. Given their diversity with respect to mechanistic details, it is quite possible that ABC transporters belonging to other subfamilies will reveal distinct conformations or intermediate states most strongly populated under turnover conditions.

## Methods

### Expression and purification of ABCG2

Human wild type ABCG2 or ABCG2 _*R184A*_ containing an N-terminal Flag-tag was expressed in HEK293-EBNA (Thermo Fisher Scientific) cells as previously described^19^. Cells were incubated at 37°C for 48-60 hours before harvesting. Harvested Cell were lysed using a Dounce homogenizer and solubilized with 1% DDM, 0.1% CHS (cholesteryl hemisuccinate) (w/v) (Anatrace), 40 mM HEPES buffer pH 7.5, 150 mM NaCl, 10% (v/v) glycerol, 1 mM PMSF (phenylmethylsulfonyl fluoride), 2 μg ml^−1^ DNaseI (Roche), and protease inhibitor cocktail (Sigma). Lysed cells were centrifuged at 100,000*g* and the supernatant was incubated with anti-Flag M2 affinity agarose gel (Sigma). ABCG2 was eluted with Flag peptide (Sigma) and applied to a Superdex 200 increase 10/300 column (GE Healthcare) in 40 mM HEPES, pH 7.5, 150 mM NaCl, 0.026% DDM and 0.0026% CHS (w/v). Peak fractions were collected for further use.

### ABCG2-nanodisc preparation

Membrane scaffold protein (MSP) 1D1 was expressed in *E. coli* and purified as described^52^. ABCG2 reconstitution in nanodiscs was performed as described^19^. In brief, BPL (brain polar lipid, Avanti Polar Lipids): CHS (cholesteryl hemisuccinate) was mixed at a 4:1 (w/w) ratio and was solubilized with a 3x molar excess of sodium cholate using a sonic bath. Lipides were then mixed with MSP1D1 and detergent -purified ABCG2 at a molar ratio of 100:5:0.2 (lipid:MSP:ABCG2). Detergent was removed by addition of Bio-Beads SM-2 (Biorad) and incubation at 4 °C overnight. Biobeads were removed and the sample was centrifuged at 100,000*g* for 30min. The supernatant was loaded on a Superdex 200 increase column and fractions containing nanodisc-reconstituted ABCG2 were collected for CryoEM grid preparation or for functional assays.

### ABCG2-liposome preparation

ABCG2-containing proteoliposomes were prepared as described^19,53^. In brief, brain polar lipid (BPL) was mixed with cholesterol (Chol) at a 4:1 (w/w) ratio. Liposomes were resuspended in transport buffer (25 mM HEPES pH 7.5, 150 mM NaCl) and extruded using a 400nm polycarbonate filter (Avanti Polar Lipids). To reconstitute ABCG2, liposomes were destabilized with 0.17% (v/v) Triton X-100. Detergent-purified ABCG2 was mixed with liposomes at a 100:1 (w/w) lipid: protein ratio. Detergent was then removed using Bio-Beads, added in multiple batches. Proteoliposomes were collected using centrifugation at 100,000*g*. The proteoliposome pellet was resuspended in transport buffer at a final lipid concentration of 10 mg ml^−1^.

### ATPase and transport assays

Both assays were performed as described previously^19,53^. ATPase assays were performed at 37 °C in the presence of 2 mM ATP and 10 mM MgCl_2_. Where indicated, E_1_S or topotecan were added at 50 μM. ATPase rates were determined using linear regression in GraphPad Prism 8. For transport assays, proteoliposomes in transport buffer (25 mM HEPES pH 7.5, 150 mM NaCl) were extruded through a 400 nm polycarbonate filter. MgCl_2_ (5 mM) and E_1_S (50 μM, containing mixtures of ^3^H-E_1_S and ^1^H-E_1_S) were added in the presence or absence of topotecan and the samples were incubated for 5 min at 30 °C. Transport reactions were initiated by adding ATP (2 mM) and stopped by adding an aliquot to ice-cold transport buffer containing unlabelled E_1_S (100 μM). Sample were filtered with a Multiscreen vacuum manifold (MSFBN6B filter plate, Millipore) and washed three times. Radioactivity trapped on the filters was measured with the microplate scintillation counter (Perkin Elmer 2450 Microbeta2). Data were analysed and curves were plotted using the nonlinear regression Michaelis–Menten analysis and initial rates were calculated using the data points from the fitting curve (GraphPad Software, La Jolla, California, USA).

### CryoEM sample preparation

All grids were prepared using a Vitrobot (FEI), with the environmental chamber set at 100% humidity and 4°C. Nanodisc-reconstituted ABCG2 (1 mg ml^-1^) was incubated with 5mM ATP, 5mM MgCl_2_, 0.5mM ADP, 100 μM topotecan or 200 μM E_1_S at room temperature for 10 min. 3.5 μl sample was applied on glow-discharged Quantifoil carbon grids (300 meshes, R 1.2/1.3 copper) immediately after incubation. Grids were blotted for 2.5s with blot force 1 and flash-frozen in a mixture of liquid ethane and propane.

### Cryo-EM data acquisition

CryoEM data of ABCG2-topotecan turnover sample was collected with a 300 keV Titan Krios (FEI) transmission electron microscope (TEM) equipped with a Gatan BioQuantum 1967 filter and a Gatan K3 camera. Images were recorded with 3 exposures per hole using EPU 2 in super--resolution mode with a 20 eV slit width of the energy filter and at a nominal magnification of 130,000 x, resulting in a calibrated super-resolution pixel size of 0.33 Å (physical pixel size 0.66 Å). Defocus was set to vary from -0.6 to −2 μm. Each image was dose fractionated to 40 frames with 1.01 s total exposure time. The dose was 1.45 e^-^/Å^2^/frame (Total dose 58 e^-^/Å^2^). The super-resolution micrographs were down-sampled twice and motion-corrected by Fourier cropping and drift-corrected and dose-weighted using MotionCor2^54^. Micrographs were visually inspected, and bad micrographs were removed manually. The topotecan turnover dataset was composed of 24,177 movies from 2 sessions.

CryoEM data of ABCG2-E_1_S turnover sample was collected on a 300 keV Titan Krios TEM equipped with a K2 Summit direct electron detector and a Quantum-LS energy filter (GIF) (20 eV zero loss filtering; Gatan Inc.). The microscope was operated with a nominal magnification of 165,000 x (the actual magnification 60,975 x). Automated data collections were performed in SerialEM with seven shots per hole, the image-shift and the coma-free setup ^55^. Dose-fractionated movies were recorded in counting mode with a physical pixel size of 0.82 Å and the defocus was set in a range from -0.8 to -2.8 μm. Each movie was dose fractionated to 40 frames with 10s total exposure time. The total dose was 50 e^-^ /Å^2^. The recorded movie stacks were initially processed by MotionCor2 (FOCUS integrated), including pre-processing procedures such as the gain-normalization, motion correction and dose weighting^56,57^. The entire dataset of ABCG2-E_1_S turnover sample from two EM sessions was composed of 15,460 movies. Cryo-EM data collection statistics in this study are presented in Supplementary table 3.

### Image processing

ABCG2-E_1_S turnover datasets from two EM sessions were processed separately with a similar procedure until 2D classifications (Supplementary Fig. 2). The aligned micrographs were manually sorted in FOCUS^56^ and then imported to CryoSPARC 2.1^31^. The contrast transfer function (CTF) and defocus values were estimated on the dose-weighted micrographs using patch-based CTF in CryoSPARC. Images with poor quality were discarded. Defocus was calibrated with a range of −0.8 to –2.8 μm. Images with a CTF-estimated resolution of better than 5.2Å were selected, resulting in a total of 11,343 micrographs. A 2D template was produced by initial 2D classifications with manually picked particles. Particles of the full dataset were auto-picked with 2D templates. After multiple rounds of 2D classification, particles were combined for 3D classification. The merged data with 645,803 particles was subjected to 3 rounds of 3D classification to classify distinct conformations. A 3D reference was generated from ab-initio 3D reconstructions. Each round was followed by a heterogenous refinement. As shown in Supplementary Fig. 2, several setting parameters were tested and applied (Round 1: 2 classes, 0.1 similarity, force hard-classification; Round 2: 2 classes, 0.8 similarity, force hard-classification; Round 2: 3 classes, 0.1 similarity, force hard-classification), resulting in a particle subset containing 221,160 particles. The best-resolved 3D class was further subjected to 3D non-uniform refinement (CryoSPARC). The map was refined with C1 symmetry by local refinement using a soft-mask. The overall resolution was 3.4Å at FSC=0.143 cutoff. A local-resolution map was generated by MonoRes^58^.

For topotecan turnover sample, the drift corrected, dose-weighted micrographs were imported in CryoSPARC v2.15^31^. Contrast transfer function (CTF) parameters were estimated with GCTF^59^. The calculated defocus parameters of micrographs were −0.3 to −2.7 μm. 1,534 particles were manually picked from selected micrographs to generate 2D template for Auto-pick. 2,623,169 particles were picked and extracted from 23,574 micrographs. After 10 rounds of classifications, 1,486,797 particles belonging to ‘good’ 2D classes were selected. Initial model was generated in Ab-Initio Reconstruction and used as reference for the first round of classification. The selected particles from 2D classification were subjected to 3D classification with 4 classes and binned 3 x. A good class from the first round of 3D classification with 750,851 particles was selected. This class was refined to 3.8 Å in Homogeneous refinement. The class was subjected to a second round of 3D classification with 3 classes and binned 3 x. A good class with 455,604 particles was selected, and it was refined to 3.45 Å in Homogeneous refinement. A third round of 3D classification was done with 455,604 particles and 3 classes. An excellent class with 311,273 particles was selected, and it was refined to 3.34 Å in Homogeneous refinement. To check if resolution can be further improved by 3D classification, a fourth round of 3D classification was done with 3 classes and binned 3 x. Two good class with 299,139 particles were selected and refined to 3.28 Å in Homogeneous refinement. The 299,139 particles subset was subjected to Global CTF Refinement followed by Local CTF refinement. Further 3D variability analysis showed there was heterogeneity in one dimension for this particle subset. To deal with the heterogeneity, a fifth round of 3D classification was performed with 3 classes and no bin. A minor class with 78,323 particles was isolated with NDB semi-closed conformation. Further homogeneous, local refinement and post-processing of the minor 3D class resulted in a 3.4 Å map named topotecan turnover-2, with automatically determined B factor of −92.8 Å^2^. The other two major classes with 220,816 particles showed NBD open conformation, thus they were combined. Further homogenous refinement, local refinement and post-processing resulted in a 3.1 Å map named topotecan turnover-1, with automatically determined B factor of −106.5 Å^2^. The final maps were refined in C1 symmetry and were directly used for modeling. Local resolution maps were generated using CryoSPARC v2.15. The data processing pipeline is shown in Supplementary Fig. 3.

3D variability analysis of topotecan turnover sample was performed with particle stacks that contributed to high-resolution 3D maps in cryoSPARC v2.15^31^. The aim was to explore heterogeneity in single particle cryo-EM data sets. A mask without lipids belt, 3 eigenvectors and 4 Å low-pass filtered were applied for 3DVA, generating simple linear “movies” of volumes. 3DVA outputs were visualized with 3D Variability Display tool^32^.

We observed both turnover-1 and turnover-2 conformations in the topotecan turnover sample, whereas we only observed the turnover-2 conformation in the E_1_S turnover sample. To avoid bias derived from reference map when doing 3D classification, we re-classified particles of the E_1_S turnover sample into 6-10 classes using turnover-1 as reference map. We expected to classify different conformations in this way. We observed some minor classes with separated NBDs. However, none of these classes produced EM maps of sufficient resolution to identify secondary structures. We can therefore exclude that the turnover-1 state with well-defined features is present in meaningful amounts in the E_1_S turnover sample. Turnover-2 is the only well-defined state of the E_1_S turnover sample.

### Model building and refinement

Coot 0.9 was used for all model building steps^60^. ABCG2-MZ29 model (PDB 6FFC) was docked into turnover maps and used as the reference for manual rebuilding^20^. The coordinate of topotecan was from ABCG2-topotecan-Fabs structure (PDB 7NEZ). E_1_S coordinate was from ABCG2-E_1_S-Fab (PDB 6HCO). Cholesterol coordinate was from ABCG2-MZ29-Fab (PDB 6HIJ) and ATP coordinate was ABCG2_E211Q_-ATP (PDB: 6HBU) ^21^. The ligands were fitted into the EM density in Coot 0.9. We generated the restraints in eLBOW^61^. Both topotecan and E_1_S turnover structures were refined against their final maps respectively in real space refinement of Phenix^62^. In the final refinement, reciprocal-space refinement of the B factors and minimization global refinements were applied together with standard geometry, rotamer, Ramachandran plot, Cβ, non-crystallographic symmetry (NCS) and secondary structure restraints. The quality of final model was assessed by MolProbity^63^. The refinement statistics are in Supplementary table 3. For model validation, we applied 0.3 Å random shifts to final models using phenix_pdb_tools^64^. The scrambled model was refined against one of the unfiltered half maps (half map A). The Fourier shell correlation between refined scrambled model and half map A was plotted as FSC_work_. The Fourier shell correlation between refined scrambled model and half map B was plotted as FSC_free_. The overlay between the FSC_work_ and FSC_free_ indicated no over-fitting presence.

### Figure preparation

Figures were prepared with PyMOL (The PyMOL Molecular Graphics System, Version 2.0 Schrödinger, LLC), GraphPad Prism 8 and UCSF ChimeraX^65^. Movies were prepared with UCSF Chimera^66^. The volume of cavity 1 in ABCG2 turnover structures was determined using POCASA^44^.

## Acknowledgments

This research was supported by the Swiss National Science Foundation through the National Centre of Competence in Research (NCCR) TransCure. We thank the Scientific Center for Optical and Electron Microscopy (ScopeM) at ETH Zürich for technical support. We thank K. Goldie, R. Irobalieva, L. Kováčik and A. Fecteau-Lefebvre for technical support with EM data collection.

## Author contributions

Q.Y. and I.M. expressed and purified wild-type ABCG2. Q.Y. cloned, expressed and purified the ABCG2_R184A_ variant. S.M.J. and I.M. reconstituted ABCG2 in lipid nanodiscs and prepared EM grids of the E_1_S turnover sample with the help of J.K. Q.Y. reconstituted ABCG2 in lipid nanodiscs and prepared EM grids of topotecan turnover sample with the help of J.K. Q.Y. reconstituted ABCG2 into liposomes and carried out all functional experiments. D.N. collected and processed cryo-EM data and determined the structure of the E_1_S turnover sample supervised by H.S. J.K. collected cryo-EM data of the topotecan turnover sample. Q.Y. processed EM data of the topotecan turnover ABCG2 sample and determined the turnover-1 and turnover-2 structures with the help of J.K. D.N. built and refined initial model of the E_1_S turnover structure. K.P.L. and Q.Y. built, refined and validated the structures. K.P.L. and Q.Y. wrote the manuscript with input from all authors. All authors contributed to revision of the manuscript.

## Figure legends

**Supplementary Figure 1. Sample preparation and cryo-EM maps a** Left, preparative SEC profile of nanodisc-reconstituted ABCG2. Right, representative SDS–PAGE analysis of main peak obtained in SEC. No reducing agent was used and ABCG2 runs as a disulfide-linked dimer. M shows marker proteins, with masses indicated on the left. The red asterisk shows the peak fraction analyzed in the gel and used for EM grid preparation. **b** Non-symmetrized cryo-EM maps of turnover states.

**Supplementary Figure 2. Cryo-EM data processing and model validation of E**_**1**_**S turnover structure. a** Representative motion-corrected 2D micrograph of E_1_S turnover sample. Black scale bar, 400 Å. **b** Representative 2D classes. **c** Flowchart of data processing of E_1_S turnover sample. Red boxes indicate classes of particles selected for the next round. **d** Local resolution of E_1_S turnover-2. **e** Map and model validation of E_1_S turnover-2. **f** Angular distribution plot for the final reconstruction of E_1_S turnover-2 from RELION.

**Supplementary Figure 3. Cryo-EM data processing and model validation of topotecan turnover structures. a** Representative motion-corrected 2D image of topotecan turnover sample. **b** Representative 2D classes. **c** Flowchart of data processing of topotecan turnover sample. Red boxes indicate classes of particles selected for the next round. **d** and **e** local resolution plot for topotecan turnover structures. **f** and **g** map and model validation of topotecan turnover structures. **h** and **i** Angular distribution plot of topotecan turnover structures from CryoSPARC.

**Supplementary Figure 4. Conformational changes in ABCG2 a** Surface representation (left) and vertical slice (right) of topotecan turnover-1 structure. Bound topotecan in cavity is shown as spheres and labeled. The leucine gate and cavity 2 are also labeled. **b** Similar to **a** but showing the topotecan turnover-2 structure. **c** Structural changes within ABCG2 protomers upon superposition of the NBDs (bound 5D3-Fab omitted for clarity). The structures are color-coded. The distinct rigid body rotations of the TMDs relative to the TMD of the topotecan-and 5D3Fab-bound structure are shown by arrows. Similarly, the rotations of the connecting helices (CnH) are indicated.

**Supplementary Figure 5. Conformational changes within TMDs** ABCG2 structures (color coding in panel **c**) are shown upon superposition of one of the TMDs (right side in panels **a** and **b**) to visualize the magnitude of the structural changes in the TMD dimer. **a** Side view of the TMDs after superposition. TM helices, coupling helices (CpH) and connecting helices (CnH) are numbered and labeled. NBDs are omitted for clarity. **b** Similar to **a** but viewed from the inside of the cell. **c** Close-up view of each TM helix in the non-superimposed TMD. Shifts at the cytoplasmic end of each TM helix or CpH or CnH are indicated.

**Supplementary Figure 6 Subdomain rotations within NBDs** Superpositions of RecA-like subdomain in NBDs of ABCG2, illustrating the rotation of the helical subdomain relative to the helical subdomain of topotecan- and Fab-bound inward-open ABCG2. The structures are color-coded and the degree of the rotation is indicated.

**Supplementary Figure 7 Biochemical characterization and transport activity of wild type ABCG2 and ABCG2**_**R184A**_**a** Preparative SEC profile of WT ABCG2 and ABCG2_R184A_ in detergent solution. Void peak (indicated by “v”) and ABCG2 dimer peak are indicated by arrows. **b** Representative SDS–PAGE analysis of ABCG2 reconstituted in proteoliposomes. Both reducing and none-reducing conditions were used. WT and R184A run as covalently (disulfide) linked dimers in non-reducing conditions and as monomers in reducing conditions. M shows marker proteins, with masses indicated on the left. **c** Initial E_1_S transport rates of wild type ABCG2 and ABCG2_R184A_ variant. Error bars show s.d. (n=3 technical replicates, same batch of liposomes).

## References

1. Krishnamurthy, P. et al. The stem cell marker Bcrp/ABCG2 enhances hypoxic cell survival through interactions with heme. J Biol Chem 279, 24218–25 (2004).

2. Kerr, I.D., Haider, A.J. & Gelissen, I.C. The ABCG family of membrane-associated transporters: you don’t have to be big to be mighty. Br J Pharmacol 164, 1767–79 (2011).

3. Enokizono, J., Kusuhara, H., Ose, A., Schinkel, A.H. & Sugiyama, Y. Quantitative investigation of the role of breast cancer resistance protein (Bcrp/Abcg2) in limiting brain and testis penetration of xenobiotic compounds. Drug Metab Dispos 36, 995–1002 (2008).

4. Maliepaard, M. et al. Subcellular localization and distribution of the breast cancer resistance protein transporter in normal human tissues. Cancer Res 61, 3458–64 (2001).

5. Matsuo, H. et al. Common defects of ABCG2, a high-capacity urate exporter, cause gout: a function-based genetic analysis in a Japanese population. Sci Transl Med 1, 5ra11 (2009).

6. Ishikawa, T., Aw, W. & Kaneko, K. Metabolic Interactions of Purine Derivatives with Human ABC Transporter ABCG2: Genetic Testing to Assess Gout Risk. Pharmaceuticals (Basel) 6, 1347–60 (2013).

7. Gottesman, M.M., Fojo, T. & Bates, S.E. Multidrug resistance in cancer: role of ATP-dependent transporters. Nat Rev Cancer 2, 48–58 (2002).

8. Szakacs, G., Paterson, J.K., Ludwig, J.A., Booth-Genthe, C. & Gottesman, M.M. Targeting multidrug resistance in cancer. Nat Rev Drug Discov 5, 219–34 (2006).

9. Morfouace, M. et al. ABCG2 Transporter Expression Impacts Group 3 Medulloblastoma Response to Chemotherapy. Cancer Res 75, 3879–89 (2015).

10. Maliepaard, M. et al. Overexpression of the BCRP/MXR/ABCP gene in a topotecan-selected ovarian tumor cell line. Cancer Res 59, 4559–63 (1999).

11. Rabindran, S.K., Ross, D.D., Doyle, L.A., Yang, W. & Greenberger, L.M. Fumitremorgin C reverses multidrug resistance in cells transfected with the breast cancer resistance protein. Cancer Res 60, 47–50 (2000).

12. Rabindran, S.K. et al. Reversal of a novel multidrug resistance mechanism in human colon carcinoma cells by fumitremorgin C. Cancer Res 58, 5850–8 (1998).

13. Allen, J.D. et al. Potent and specific inhibition of the breast cancer resistance protein multidrug transporter in vitro and in mouse intestine by a novel analogue of fumitremorgin C. Mol Cancer Ther 1, 417–25 (2002).

14. Puentes, C.O. et al. Solid phase synthesis of tariquidar-related modulators of ABC transporters preferring breast cancer resistance protein (ABCG2). Bioorg Med Chem Lett 21, 3654–7 (2011).

15. Pick, A., Klinkhammer, W. & Wiese, M. Specific inhibitors of the breast cancer resistance protein (BCRP). ChemMedChem 5, 1498–505 (2010).

16. Kohler, S.C. & Wiese, M. HM30181 Derivatives as Novel Potent and Selective Inhibitors of the Breast Cancer Resistance Protein (BCRP/ABCG2). Journal of Medicinal Chemistry 58, 3910–3921 (2015).

17. Miyata, H. et al. Identification of Febuxostat as a New Strong ABCG2 Inhibitor: Potential Applications and Risks in Clinical Situations. Frontiers in Pharmacology 7(2016).

18. Weidner, L.D. et al. The Inhibitor Ko143 Is Not Specific for ABCG2. J Pharmacol Exp Ther 354, 384–93 (2015).

19. Taylor, N.M.I. et al. Structure of the human multidrug transporter ABCG2. Nature 546, 504–509 (2017).

20. Jackson, S.M. et al. Structural basis of small-molecule inhibition of human multidrug transporter ABCG2. Nature Structural & Molecular Biology 25, 333-+ (2018).

21. Manolaridis, I. et al. Cryo-EM structures of a human ABCG2 mutant trapped in ATP-bound and substrate-bound states. Nature 563, 426–430 (2018).

22. Orlando, B.J. & Liao, M. ABCG2 transports anticancer drugs via a closed-to-open switch. Nat Commun 11, 2264 (2020).

23. Kowal, J. et al. Structural basis of drug recognition by the multidrug transporter ABCG2. bioRxiv (2021).

24. Gose, T. et al. ABCG2 requires a single aromatic amino acid to “clamp” substrates and inhibitors into the binding pocket. FASEB J 34, 4890–4903 (2020).

25. Zhou, S. et al. The ABC transporter Bcrp1/ABCG2 is expressed in a wide variety of stem cells and is a molecular determinant of the side-population phenotype. Nat Med 7, 1028–34 (2001).

26. Dawson, R.J. & Locher, K.P. Structure of a bacterial multidrug ABC transporter. Nature 443, 180–5 (2006).

27. Kim, Y. & Chen, J. Molecular structure of human P-glycoprotein in the ATP-bound, outward-facing conformation. Science 359, 915–919 (2018).

28. Korkhov, V.M., Mireku, S.A. & Locher, K.P. Structure of AMP-PNP-bound vitamin B12 transporter BtuCD-F. Nature 490, 367–72 (2012).

29. Alam, A. et al. Structure of a zosuquidar and UIC2-bound human-mouse chimeric ABCB1. Proc Natl Acad Sci U S A 115, E1973–E1982 (2018).

30. Frank, G.A. et al. Cryo-EM Analysis of the Conformational Landscape of Human P-glycoprotein (ABCB1) During its Catalytic Cycle. Mol Pharmacol 90, 35–41 (2016).

31. Punjani, A., Rubinstein, J.L., Fleet, D.J. & Brubaker, M.A. cryoSPARC: algorithms for rapid unsupervised cryo-EM structure determination. Nat Methods 14, 290–296 (2017).

32. Punjani, A. & Fleet, D.J. 3D Variability Analysis: Resolving continuous flexibility and discrete heterogeneity from single particle cryo-EM. J Struct Biol, 107702 (2021).

33. Beis, K. Structural basis for the mechanism of ABC transporters. Biochem Soc Trans 43, 889–93 (2015).

34. Locher, K.P. Mechanistic diversity in ATP-binding cassette (ABC) transporters. Nat Struct Mol Biol 23, 487–93 (2016).

35. Rees, D.C., Johnson, E. & Lewinson, O. ABC transporters: the power to change. Nat Rev Mol Cell Biol 10, 218–27 (2009).

36. Hollenstein, K., Frei, D.C. & Locher, K.P. Structure of an ABC transporter in complex with its binding protein. Nature 446, 213–6 (2007).

37. Miyake, K. et al. Molecular cloning of cDNAs which are highly overexpressed in mitoxantrone-resistant cells: Demonstration of homology to ABC transport genes. Cancer Research 59, 8–13 (1999).

38. Doyle, L.A. et al. A multidrug resistance transporter from human MCF-7 breast cancer cells. Proc Natl Acad Sci U S A 95, 15665–70 (1998).

39. Robey, R.W. et al. Mutations at amino-acid 482 in the ABCG2 gene affect substrate and antagonist specificity. British Journal of Cancer 89, 1971–1978 (2003).

40. Honjo, Y. et al. Acquired mutations in the MXR/BCRP/ABCP gene alter substrate specificity in MXR/BCRP/ABCP-overexpressing cells. Cancer Res 61, 6635–9 (2001).

41. Komatani, H. et al. Identification of breast cancer resistant protein/mitoxantrone resistance/placenta-specific, ATP-binding cassette transporter as a transporter of NB-506 and J-107088, topoisomerase I inhibitors with an indolocarbazole structure. Cancer Research 61, 2827–2832 (2001).

42. Pozza, A., Perez-Victoria, J.M., Sardo, A., Ahmed-Belkacem, A. & Di Pietro, A. Purification of breast cancer resistance protein ABCG2 and role of arginine-482. Cellular and Molecular Life Sciences 63, 1912–1922 (2006).

43. Ejendal, K.F., Diop, N.K., Schweiger, L.C. & Hrycyna, C.A. The nature of amino acid 482 of human ABCG2 affects substrate transport and ATP hydrolysis but not substrate binding. Protein Sci 15, 1597–607 (2006).

44. Yu, J., Zhou, Y., Tanaka, I. & Yao, M. Roll: a new algorithm for the detection of protein pockets and cavities with a rolling probe sphere. Bioinformatics 26, 46–52 (2010).

45. Alam, A., Kowal, J., Broude, E., Roninson, I. & Locher, K.P. Structural insight into substrate and inhibitor discrimination by human P-glycoprotein. Science 363, 753-+ (2019).

46. Nosol, K. et al. Cryo-EM structures reveal distinct mechanisms of inhibition of the human multidrug transporter ABCB1. Proc Natl Acad Sci U S A 117, 26245–26253 (2020).

47. Wang, L. et al. Characterization of the kinetic cycle of an ABC transporter by single-molecule and cryo-EM analyses. Elife 9(2020).

48. Hofmann, S. et al. Conformation space of a heterodimeric ABC exporter under turnover conditions. Nature 571, 580–583 (2019).

49. Johnson, Z.L. & Chen, J. ATP Binding Enables Substrate Release from Multidrug Resistance Protein 1. Cell 172, 81–89 e10 (2018).

50. Shintre, C.A. et al. Structures of ABCB10, a human ATP-binding cassette transporter in apo- and nucleotide-bound states. Proc Natl Acad Sci U S A 110, 9710–5 (2013).

51. Ambudkar, S.V., Kim, I.W., Xia, D. & Sauna, Z.E. The A-loop, a novel conserved aromatic acid subdomain upstream of the Walker A motif in ABC transporters, is critical for ATP binding. FEBS Lett 580, 1049–55 (2006).

52. Ritchie, T.K. et al. Chapter 11 - Reconstitution of membrane proteins in phospholipid bilayer nanodiscs. Methods Enzymol 464, 211–31 (2009).

53. Geertsma, E.R., Mahmood, N.A.B.N., Schuurman-Wolters, G.K. & Poolman, B. Membrane reconstitution of ABC transporters and assays of translocator function. Nature Protocols 3, 256–266 (2008).

54. Li, X.M. et al. Electron counting and beam-induced motion correction enable near-atomic-resolution single-particle cryo-EM. Nature Methods 10, 584-+ (2013).

55. Mastronarde, D.N. SerialEM: A Program for Automated Tilt Series Acquisition on Tecnai Microscopes Using Prediction of Specimen Position. Microscopy and Microanalysis 9, 1182–1183 (2003).

56. Biyani, N. et al. Focus: The interface between data collection and data processing in cryo-EM. J Struct Biol 198, 124–133 (2017).

57. Li, X. et al. Electron counting and beam-induced motion correction enable near-atomic-resolution single-particle cryo-EM. Nat Methods 10, 584–90 (2013).

58. Vilas, J.L. et al. MonoRes: Automatic and Accurate Estimation of Local Resolution for Electron Microscopy Maps. Structure 26, 337–344 e4 (2018).

59. Zhang, K. Gctf: Real-time CTF determination and correction. J Struct Biol 193, 1–12 (2016).

60. Emsley, P., Lohkamp, B., Scott, W.G. & Cowtan, K. Features and development of Coot. Acta Crystallographica Section D-Biological Crystallography 66, 486–501 (2010).

61. Moriarty, N.W., Grosse-Kunstleve, R.W. & Adams, P.D. electronic Ligand Builder and Optimization Workbench (eLBOW): a tool for ligand coordinate and restraint generation. Acta Crystallogr D Biol Crystallogr 65, 1074–80 (2009).

62. Afonine, P.V. et al. Real-space refinement in PHENIX for cryo-EM and crystallography. Acta Crystallogr D Struct Biol 74, 531–544 (2018).

63. Chen, V.B. et al. MolProbity: all-atom structure validation for macromolecular crystallography. Acta Crystallogr D Biol Crystallogr 66, 12–21 (2010).

64. Adams, P.D. et al. PHENIX: a comprehensive Python-based system for macromolecular structure solution. Acta Crystallogr D Biol Crystallogr 66, 213–21 (2010).

65. Goddard, T.D. et al. UCSF ChimeraX: Meeting modern challenges in visualization and analysis. Protein Sci 27, 14–25 (2018).

66. Pettersen, E.F. et al. UCSF Chimera--a visualization system for exploratory research and analysis. J Comput Chem 25, 1605–12 (2004).

