## Supplemental Tables and Figures for "Structures of ABCG2 under turnover conditions reveal a key step in drug transport mechanism"

| Structure | topotecan- and 5D3-Fab-bound inward-open conformation (PDB: 7NEZ) | topotecan turnover-1 | $E_1S$ turnover-2 | Closed conformation (PDB: 6HBU) |
|---|---|---|---|---|
| RMSD relative to topotecan turnover-2 conformation | 2.729 Å | 2.162 Å | 0.504 Å | 3.351 Å |

Supplementary table 1. Overall R.M.S.D between ABCG2 structures of different conformations

| **Substrate** | **$EC_{50}$ (μM) (ATPase)** |
|---|---|
| Mitoxantrone | 18.3 ± 2.2[a] |
| Topotecan | 11.7 ± 0.3[a] |
| Tariquidar | 0.08 ± 0.01[a] |
| $E_1S$ | 15.7±0.9 [b] |
| $E_1S$+ Fab | 18.0±4.7[b] |

Supplementary table 2. $EC_{50}$ of ATPase stimulation of ABCG2 by substrates. The data is combined from previous study to allow direct comparison.
[a]Data taken from[23]
[b]Data taken from[19]

| Cryo-EM data collection, refinement and validation statistics | | | |
|---|---|---|---|
| Data collection and processing | $E_1S$ turnover | Topotecan turnover | |
| Map name | turnover-2 | turnover-1 | turnover-2 |
| Energy filters & detector | Quantum-LS / K2 | BioQuantum / K3 | |
| Energy window (eV) | 20 | 20 | |
| Magnification (nominal) | 165,000 x | 130,000 x | |
| Voltage (kV) | 300 | 300 | |
| Electron exposure per movie ($e^-/Å^2$) | 50 | 58 | |
| Electron exposure per frame ($e^-/Å^2$/frame) | 1.25 | 1.45 | |
| Defocus range (μm) | 0.8-2.8 | 0.6-2.0 | |
| Pixel size (Å) | 0.82 | 0.66 | |
| symmetry | C1 | C1 | |
| Initial particle images (No.) | 645,803 | 2,623,169 | |
| Final particle images (No.) | 221,160 | 220,816 | 78,323 |
| Map resolution (Å) | 3.40 | 3.13 | 3.34 |
| FSC threshold | 0.143 | 0.143 | 0.143 |
| Map resolution range (Å) | 2.7-20.0 | 2.65-20 | 3-20 |
| **Model refinement** | | | |
| Initial model used | 6HCO | 6HCO | 6HCO |
| Model resolution(Å) | 3.8 | 3.2 | 3.5 |
| FSC threshold | 0.5 | 0.5 | 0.5 |
| model resolution range (Å) | 3.8-30 | 3.2-254 | 3.5-254 |
| Map sharpening B factor ($Å^2$) | -118 | -106.5 | 92.8 |
| Model composition | | | |
| Non-hydrogen atoms | 9,148 | 8,999 | 9,195 |
| protein residues | 1,150 | 1,138 | 1,144 |
| Ligands | ATP: 2 CLR:4 $E_1S$:1 | ATP: 2 CLR:2 Topotecan:1 | ATP: 2 CLR: 4 Topotecan:1 PLC: 2 |
| B factors ($Å^2$) | | | |
| protein | 65.86/155.94/96.41 | 17.21/65.49/35.70 | 45.05/100.99/64.77 |
| ligand | 45.56/118.32/83.62 | 18.42/48.88/31.46 | 45.56/77.34/62.38 |
| R.m.s.d deviations | | | |
| Bond lengths (Å) | 0.008 | 0.006 | 0.009 |
| bond angles (°) | 0.875 | 0.655 | 0.657 |
| Validation | | | |
| MolProbity score | 1.75 | 1.13 | 1.54 |
| Clash score | 6.54 | 2.04 | 4.13 |
| Poor rotamers (%) | 0.41 | 0.52 | 0.41 |
| Ramachandran plot | | | |
| Favored (%) | 94.18 | 97.15 | 95.04 |
| Allowed (%) | 5.29 | 2.85 | 4.79 |
| Disallowed (%) | 0.53 | 0 | 0.18 |

Supplementary table 3. Cryo-EM data collection, refinement and validation statistics

Suppl. Fig. 1

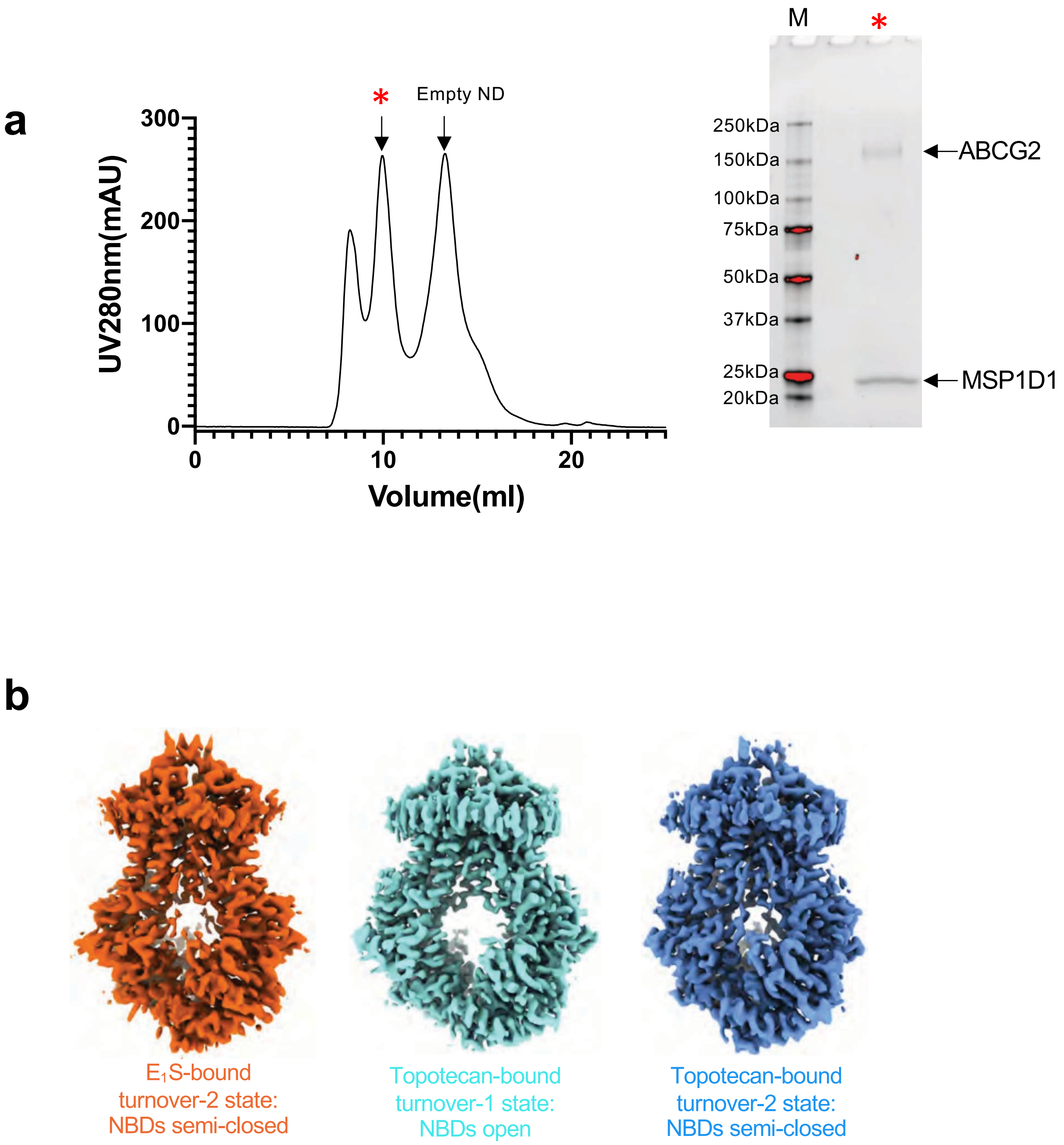
a
*
Empty ND
300
200
100
0
UV280nm(mAU)
0
10
20
Volume(ml)
M
*
250kDa
150kDa
100kDa
75kDa
50kDa
37kDa
25kDa
20kDa
ABCG2
MSP1D1
b
E1S-bound turnover-2 state: NBDs semi-closed
Topotecan-bound turnover-1 state: NBDs open
Topotecan-bound turnover-2 state: NBDs semi-closed

### Suppl. Fig. 2

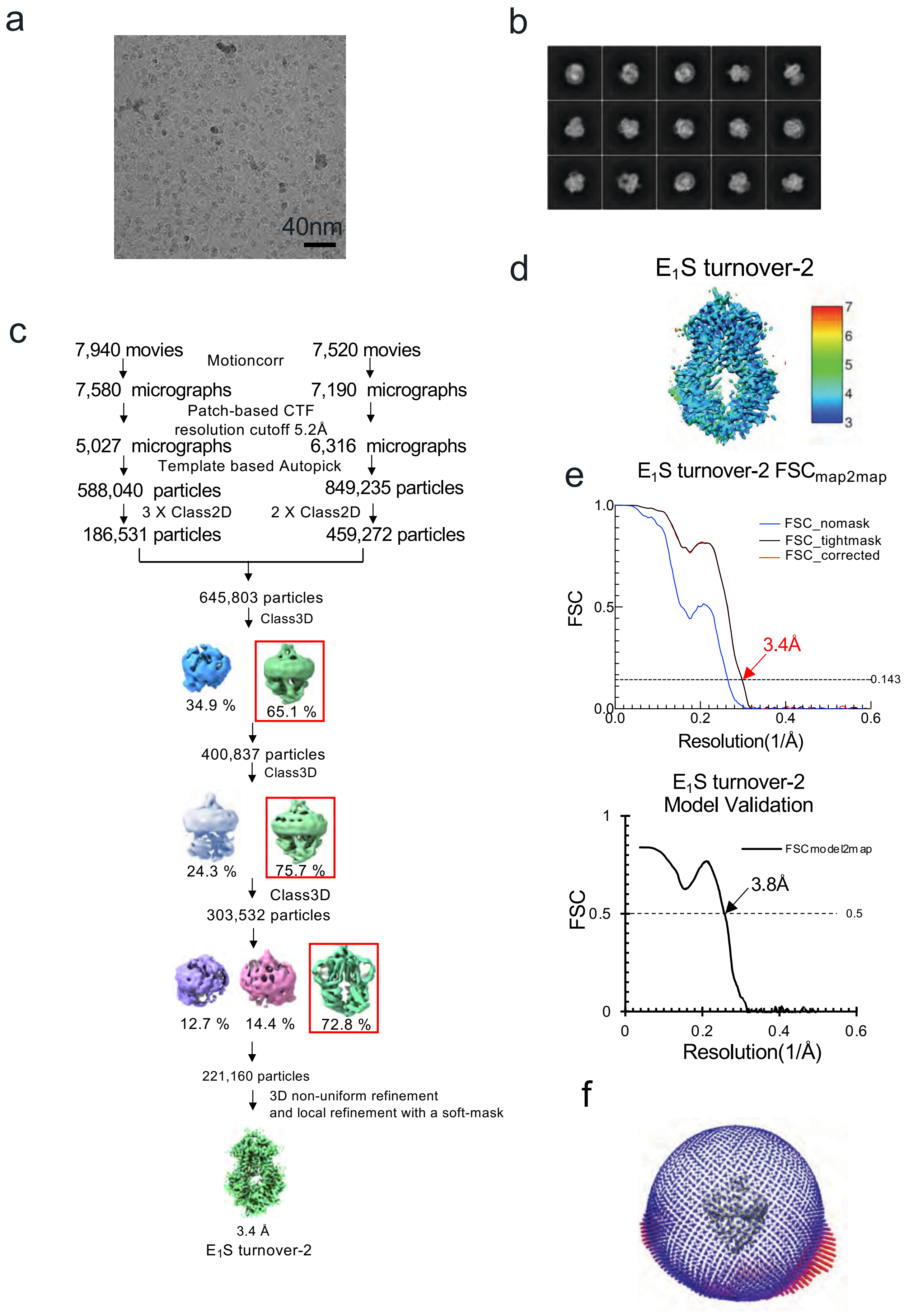
a
40nm
b
c
7,940 movies
Motioncorr
7,520 movies
7,580 micrographs
7,190 micrographs
Patch-based CTF
resolution cutoff 5.2Å
5,027 micrographs
6,316 micrographs
Template based Autopick
588,040 particles
849,235 particles
3 X Class2D
2 X Class2D
186,531 particles
459,272 particles
645,803 particles
Class3D
34.9 %
65.1 %
400,837 particles
Class3D
24.3 %
75.7 %
Class3D
303,532 particles
12.7 %
14.4 %
72.8 %
221,160 particles
3D non-uniform refinement
and local refinement with a soft-mask
3.4 Å
$E_1S$ turnover-2
d
$E_1S$ turnover-2
7
6
5
4
3
e
$E_1S$ turnover-2 $FSC_{map2map}$
FSC_nomask
FSC_tightmask
FSC_corrected
FSC
1.0
0.5
0.0
3.4Å
0.143
0.0
0.2
0.4
0.6
Resolution(1/Å)
$E_1S$ turnover-2
Model Validation
FSCmodel2map
1
FSC
0.5
0
3.8Å
0.5
0
0.2
0.4
0.6
Resolution(1/Å)
f

### Suppl. Fig. 3

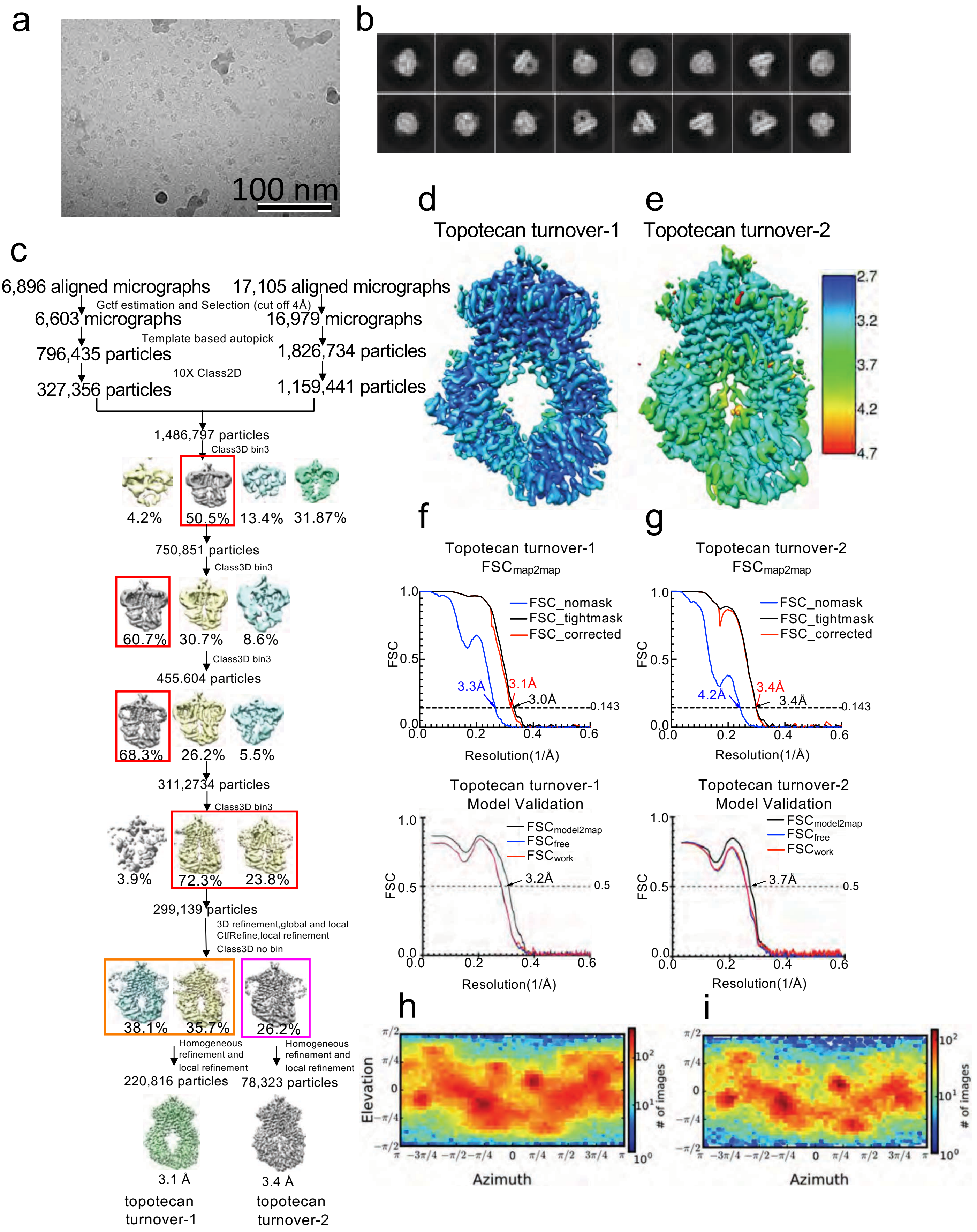
a
100 nm
b
c
6,896 aligned micrographs
17,105 aligned micrographs
Gctf estimation and Selection (cut off 4Å)
6,603 micrographs
16,979 micrographs
Template based autopick
796,435 particles
1,826,734 particles
10X Class2D
327,356 particles
1,159,441 particles
1,486,797 particles
Class3D bin3
4.2%
50.5%
13.4%
31.87%
750,851 particles
Class3D bin3
60.7%
30.7%
8.6%
Class3D bin3
455.604 particles
68.3%
26.2%
5.5%
311,2734 particles
Class3D bin3
3.9%
72.3%
23.8%
299,139 particles
3D refinement,global and local
CtfRefine,local refinement
Class3D no bin
38.1%
35.7%
26.2%
Homogeneous refinement and local refinement
Homogeneous refinement and local refinement
220,816 particles
78,323 particles
3.1 Å
3.4 Å
topotecan turnover-1
topotecan turnover-2
d
Topotecan turnover-1
e
Topotecan turnover-2
2.7
3.2
3.7
4.2
4.7
f
Topotecan turnover-1
FSCmap2map
FSC_nomask
FSC_tightmask
FSC_corrected
3.3Å
3.1Å
3.0Å
0.143
FSC
Resolution(1/Å)
g
Topotecan turnover-2
FSCmap2map
FSC_nomask
FSC_tightmask
FSC_corrected
4.2Å
3.4Å
3.4Å
0.143
FSC
Resolution(1/Å)
Topotecan turnover-1
Model Validation
FSCmodel2map
FSCfree
FSCwork
3.2Å
0.5
FSC
Resolution(1/Å)
Topotecan turnover-2
Model Validation
FSCmodel2map
FSCfree
FSCwork
3.7Å
0.5
FSC
Resolution(1/Å)
h
Elevation
Azimuth
### of images
i
Azimuth
### of images

Suppl. Fig. 4

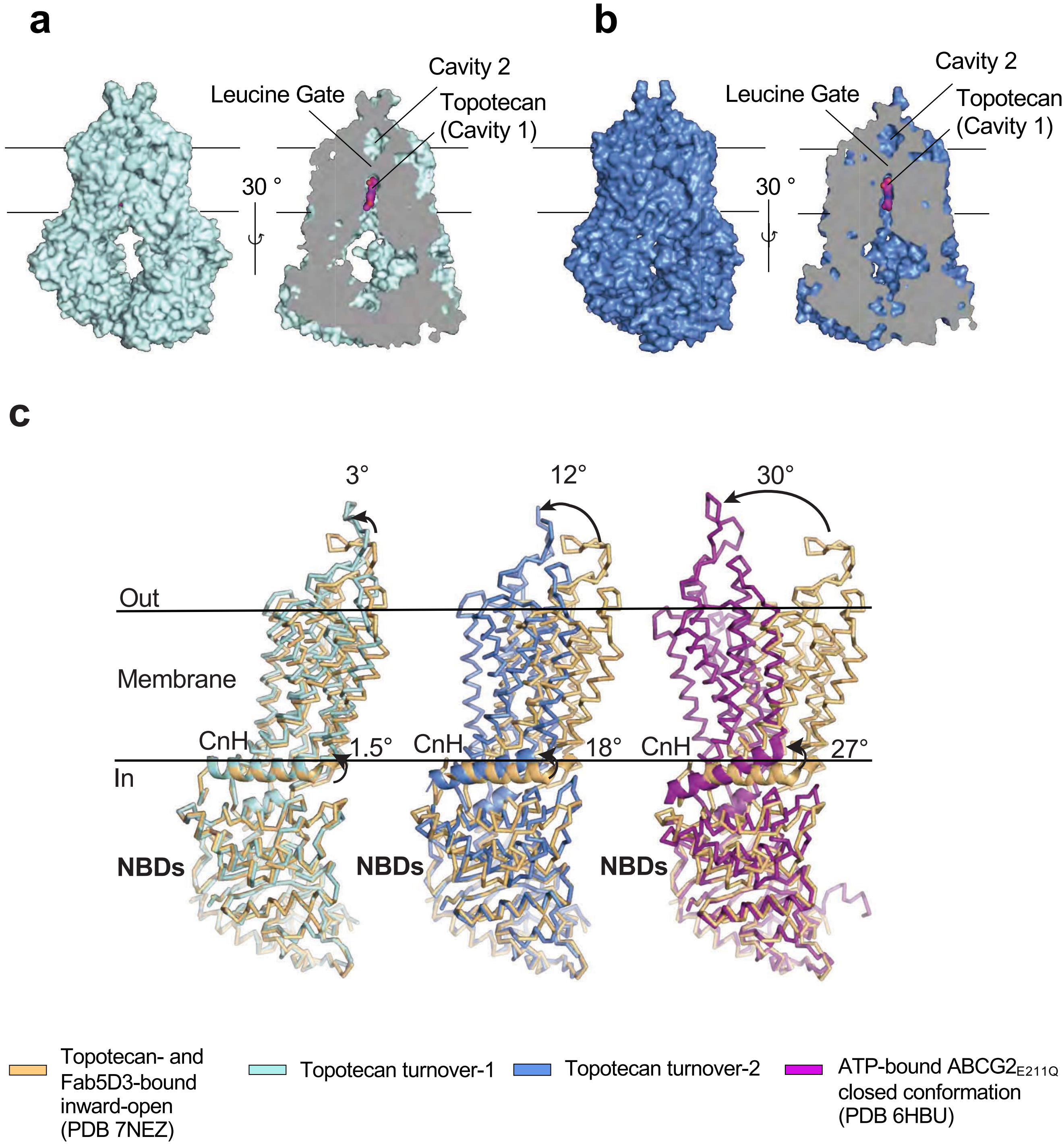
a
b
Cavity 2
Leucine Gate
Topotecan
(Cavity 1)
30 °
c
3°
12°
30°
Out
Membrane
CnH
1.5°
18°
27°
In
NBDs
Topotecan- and Fab5D3-bound inward-open (PDB 7NEZ)
Topotecan turnover-1
Topotecan turnover-2
ATP-bound ABCG2E211Q closed conformation (PDB 6HBU)

Suppl. Fig. 5

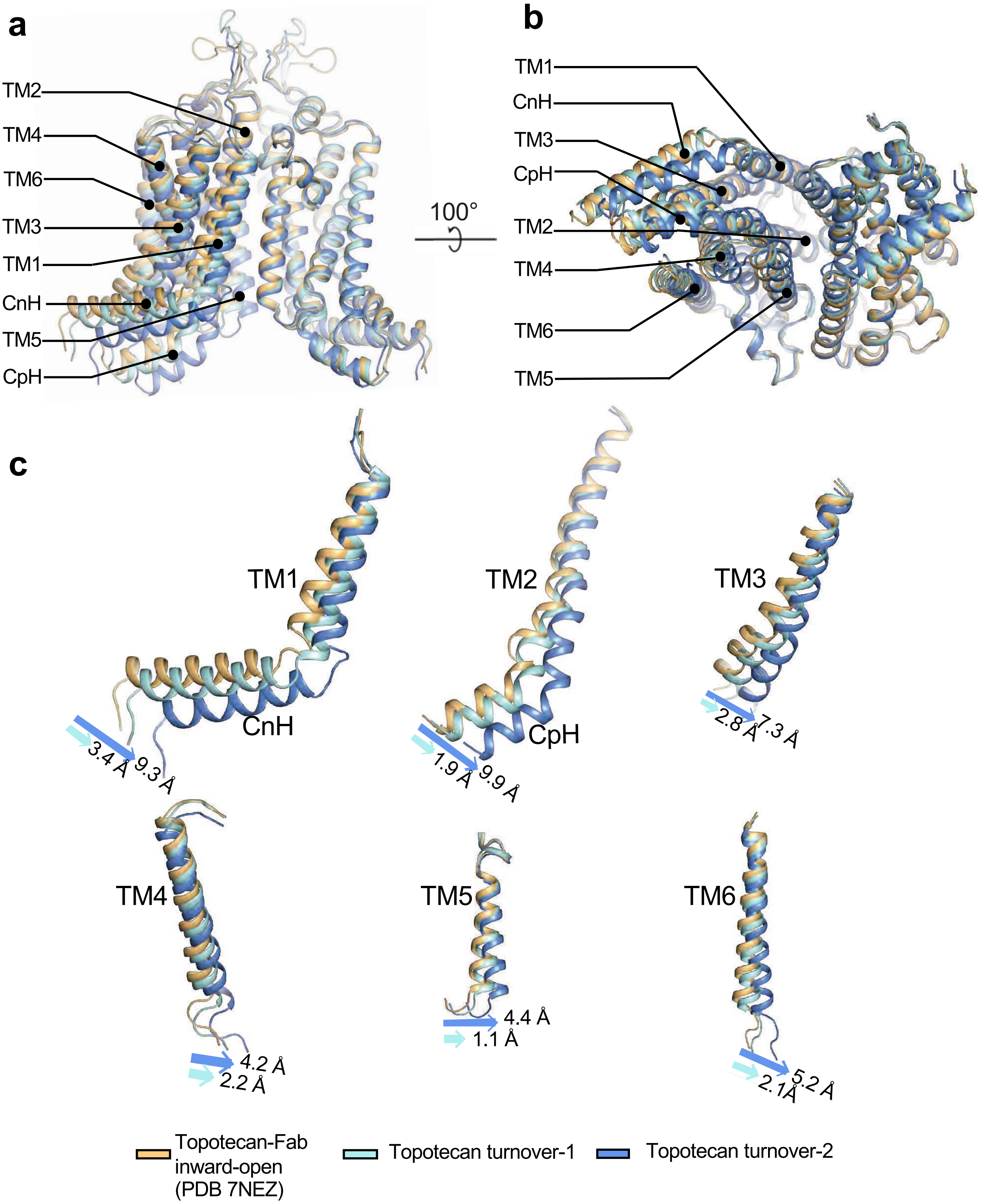

a
TM2
TM4
TM6
TM3
TM1
CnH
TM5
CpH
100°
b
TM1
CnH
TM3
CpH
TM2
TM4
TM6
TM5
c
TM1
CnH
3.4 Å
9.3 Å
TM2
CpH
1.9 Å
9.9 Å
TM3
2.8 Å
7.3 Å
TM4
4.2 Å
2.2 Å
TM5
4.4 Å
1.1 Å
TM6
5.2 Å
2.1Å
Topotecan-Fab inward-open (PDB 7NEZ)
Topotecan turnover-1
Topotecan turnover-2

### Suppl. Fig. 6

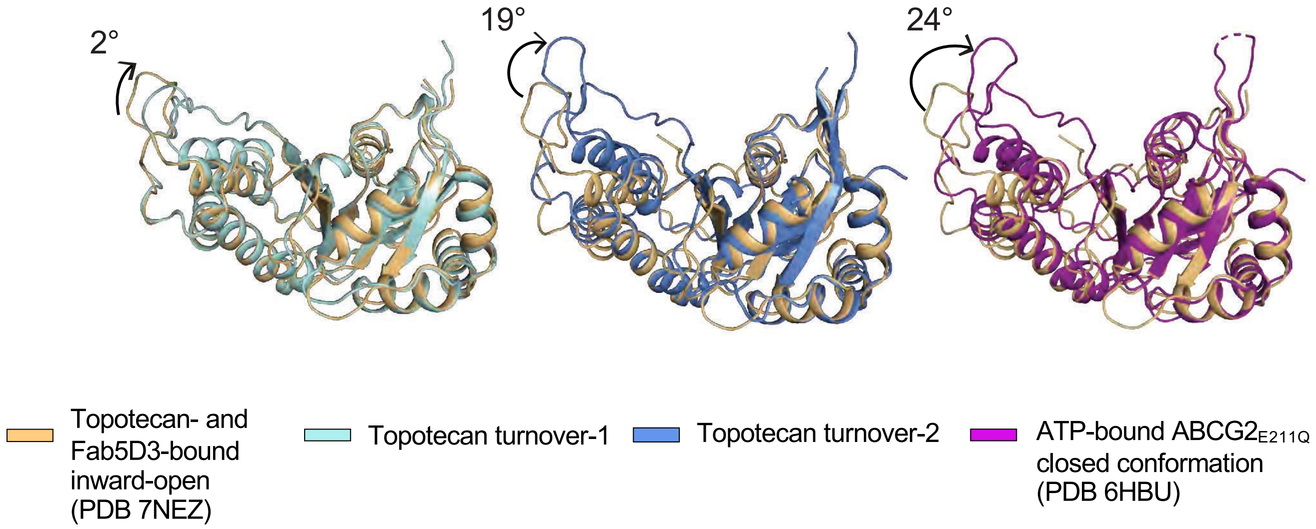

2°
19°
24°
Topotecan- and Fab5D3-bound inward-open (PDB 7NEZ)
Topotecan turnover-1
Topotecan turnover-2
ATP-bound ABCG2$_{E211Q}$ closed conformation (PDB 6HBU)

Suppl. Fig. 7

**a**

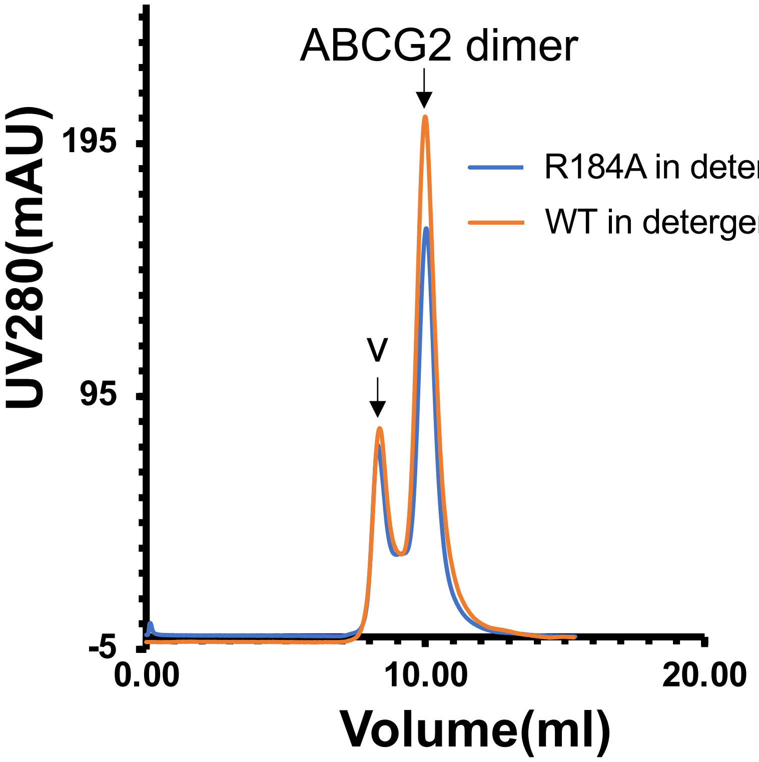


**b**

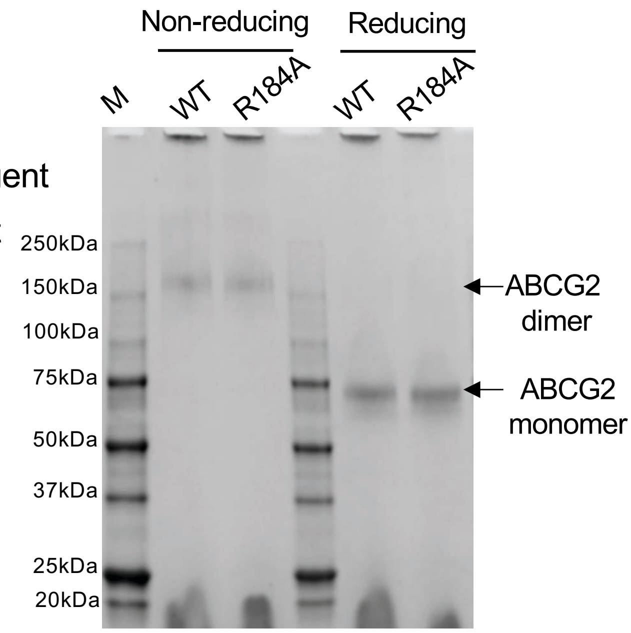


**c**

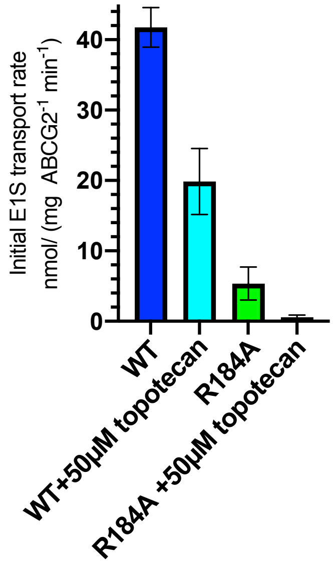


| **Sample** | **Initial rate (mol $E_1S$/mol G2 dimer/s)** |
|---|---|
| ABCG2 WT | 0.1006 ± 0.0117 |
| ABCG2 WT+50μM topotecan | 0.0479 ± 0.0196 |
| ABCG2 R184A | 0.0129 ± 0.0097 |
| ABCG2 R184A +50μM topotecan | - |
